# Quantifying the impact of electric fields on single-cell motility

**DOI:** 10.1101/2021.01.22.427762

**Authors:** TP Prescott, K Zhu, M Zhao, RE Baker

## Abstract

Cell motility in response to environmental cues forms the basis of many developmental processes in multicellular organisms. One such environmental cue is an electric field (EF), which induces a form of motility known as electrotaxis. Electrotaxis has evolved in a number of cell types to guide wound healing, and has been associated with different cellular processes, suggesting that observed electrotactic behaviour is likely a combination of multiple distinct effects arising from the presence of an EF. In order to determine the different mechanisms by which observed electrotactic behaviour emerges, and thus to design EFs that can be applied to direct and control electrotaxis, researchers require accurate quantitative predictions of cellular responses to externally-applied fields. Here, we use mathematical modelling to formulate and parametrise a variety of hypothetical descriptions of how cell motility may change in response to an EF. We calibrate our model to observed data using synthetic likelihoods and Bayesian sequential learning techniques, and demonstrate that EFs bias cellular motility through only one of a selection of hypothetical mechanisms. We also demonstrate how the model allows us to make predictions about cellular motility under different EFs. The resulting model and calibration methodology will thus form the basis for future data-driven and model-based feedback control strategies based on electric actuation.

**SIGNIFICANCE:** Electrotaxis is attracting much interest and development as a technique to control cell migration due to the precision of electric fields as actuation signals. However, precise control of electrotactic migration relies on an accurate model of how cell motility changes in response to applied electric fields. We present and calibrate a parametrised stochastic model that accurately replicates experimental single-cell data and enables the prediction of input–output behaviour while quantifying uncertainty and stochasticity. The model allows us to elucidate and quantify how electric fields perturb the motile behaviour of the cell. This model and the associated simulation-based calibration methodology will be central to future developments in the control of electrotaxis.

## INTRODUCTION

Cell migration underpins key physiological processes central to developmental biology, as well as wound healing and tissue regeneration, and it plays a crucial role in invasive, metastatic cancers. There are ongoing efforts to intervene in and influence these phenomena to, for example, inhibit metastasis (1) or accelerate wound healing (2). However, the cellular processes driving collective migration are complex and multifaceted, deriving from diverse physical mechanisms and various external stimuli (3), making it challenging for researchers to accurately and robustly direct cell motility. Due to the ease with which electric fields can be controlled and applied to cells, research into the control of cell motility has recently focused on exploiting *electrotaxis* (also known as galvanotaxis) (3–5). However, the precise effects of electric fields on intracellular processes and thus on cell motility are not fully understood, making quantitative predictions and control policy design impractical.

Electrotactic cells have been observed to change their motile behaviour in response to the presence of a direct current (DC) electric field (EF) (3–7). Researchers seeking to control cell motility exploit this phenomenon by applying external electrical cues to cell populations (2, 4–9). The key advantages of using electrical cues to guide cell migration include the ability to exploit endogenous, evolved biological functionality to respond to precisely controllable DC EFs. This compares favourably to using chemoattractants to guide motility, since chemical signals experienced by the cell cannot be so precisely or flexibly controlled, especially dynamically, and chemoattractants are usually highly cell-specific. In contrast, light-directed motility allows for precise actuation signals. However, it requires sophisticated optogenetic manipulations of the cell population under control (10). As such, EFs provide a relatively precise and simply implemented actuation signal to achieve specified motile behaviours.

While an important strength of electrotactic cell control is that applying an EF for actuation is flexible enough to apply to any electrotactic cell type, the precise signal to be applied in order to achieve any specified goal needs to be carefully calibrated. At the most basic level, even the direction of migration within the same DC field has been shown to vary across different cell types, and within one cell type under different experimental conditions (5, 11). More broadly, a large number of biochemical and biophysical mechanisms have been implicated in the electrotactic response across different cell types (3). Each electrotactic mechanism, which may co-exist in combination at unknown relative strengths, may induce distinct observable effects on the dynamics of cellular motility. Overcoming this uncertainty in the observable electrotactic response is a fundamental challenge for designing EFs to control cell motility.

Mathematical models are a vital tool for quantifying the different ways in which cells can change their motility in response to EFs (12–14). In this paper we describe a parametrised stochastic model of the motile behaviour of a single human corneal epithelial cell, in which the cell’s motility is driven by an internal polarity, in combination with the external influence of a DC EF. We assume that the cell can undergo both spontaneous and electrotactic polarisation. The model allows us to describe mathematically four distinct ways an EF may influence motility. We use experimentally observed trajectories of single cells, both with and without applied EFs, to calibrate the parameters of this model, thereby quantifying the extent to which different aspects of cell motility are impacted by the EF. The resulting calibrated model provides a vital first step towards being able to design feedback control policies and provide robustness guarantees, which are necessary if electrotaxis is to be used to control cell motility in practical applications such as wound healing or tissue engineering.

### Single-cell modelling

The agent-based modelling framework used in this work follows standard modelling assumptions outlined in (13). Specifically, we model the evolution of the velocity of a single cell in the overdamped regime, so that cell velocity is proportional to the sum of non-frictional forces on the cell. We provide full details on the mathematical model in *Materials and Methods* and in the *Supplementary Material*.

In the absence of any EF, the only non-frictional force acting on the cell is assumed to be an active force arising from the internal *polarity* of the cell. Thus, the cell velocity, **v** = **v**_cell_, is comprised of a single component. A preliminary analysis of single-cell motility data, described more fully in the *Supplementary Material*, suggests that the cell velocity arises from a cell being *polarised* in a particular direction, and that the direction of polarisation drifts stochastically over time. A polarised cell has a positive speed, parametrised by a modal value, ||**v**_cell_|| ≈ *v*, where the scalar-valued parameter *v* > 0 has dimensions μm min^−1^. In addition to random changes in cell speed, preliminary analysis also suggests that the direction of cell motion stochastically evolves according to a persistent random walk, such that the autocorrelation between displacement directions decays as the time lag increases. Thus, the direction of cell motion (in the absence of an EF) is assumed to vary according to an unbiased random walk with positive timescale constant *D* > 0, with dimensions min^−1^, which characterises the rate of decay in the autocorrelation of the polarisation direction over time. Eq. (2) in *Materials and Methods* provides the mathematical formulation of this model.

We hypothesise that a vector-valued DC EF, **u**, can affect cell motility in a variety of ways. We use a number of extensions of the model in order to implement different ways in which motility may be impacted by the EF, specifying, in particular, four distinct ways in which it may affect the dynamics of a motile cell. We parametrise the magnitude of each hypothesised electrotactic effect, observed at a reference EF strength of 200 mV mm^−1^, by the parameters *γ*_1_, *γ*_2_, *γ*_3_ and *γ*_4_, such that if *γ*_i_ = 0 then the corresponding hypothesised effect is not included in the model. Eq. (3) in *Materials and Methods* provides the mathematical formulation of this model.

The four means by which we model cell motility to be perturbed by the EF are:

**Velocity bias** (*γ*_1_) The EF imparts an additional component of force on the cell. The resulting velocity, **v** = **v**_cell_ + **v**_EF_, is thus the sum of two components: the original polarity component, **v**_cell_, and an EF component, **v**_EF_. The EF velocity component acts in the direction of the field with magnitude *γ*_1_*v*.
**Speed increase** (*γ*_2_) Polarised cells travel more quickly under the influence of an EF in the direction in which they are polarised. The modal magnitude of **v**_cell_ for polarised cells is increased by *γ*_2_*v*.
**Speed alignment** (*γ*_3_) Polarised cells travel more quickly when the direction of their polarisation aligns with the EF, but slower if opposed to the EF. The modal magnitude of **v**_cell_ for polarised cells is increased by *γ*_3_*v* cos(*θ*), where *θ* is the angle between **v**_cell_ (i.e. the polarity direction) and the EF direction.
**Polarity bias** (*γ*_4_) The random walk determining cell polarity is biased so that cells preferentially polarise in the direction of the EF. The strength of this bias is parametrised by *γ*_4_.

Two models can be distinguished: the *autonomous* model, where no EF is applied, and the *electrotactic* model, where a reference strength EF is applied. In each of these models, the cell velocity at time *t*, denoted **v**(*t*), undergoes a random walk. Figure 1 characterises each of these models by depicting the stationary probability distribution of this random walk. The top plot shows that without an applied EF the modal cell speed is near *v*, with direction chosen uniformly at random. The bottom plot of this figure demonstrates how each electrotactic effect, quantified by the value of *γ_i_* for *i* = 1, 2, 3, 4, can be interpreted in terms of the probability distribution of the cell velocity: *γ*_1_ translates the velocity distribution uniformly in the direction of the field; *γ*_2_ rescales the domain of the distribution; *γ*_3_ parametrises asymmetry in the shape of the velocity distribution; and *γ*_4_ parametrises asymmetry in the density of the velocity distribution.

**Figure 1:**
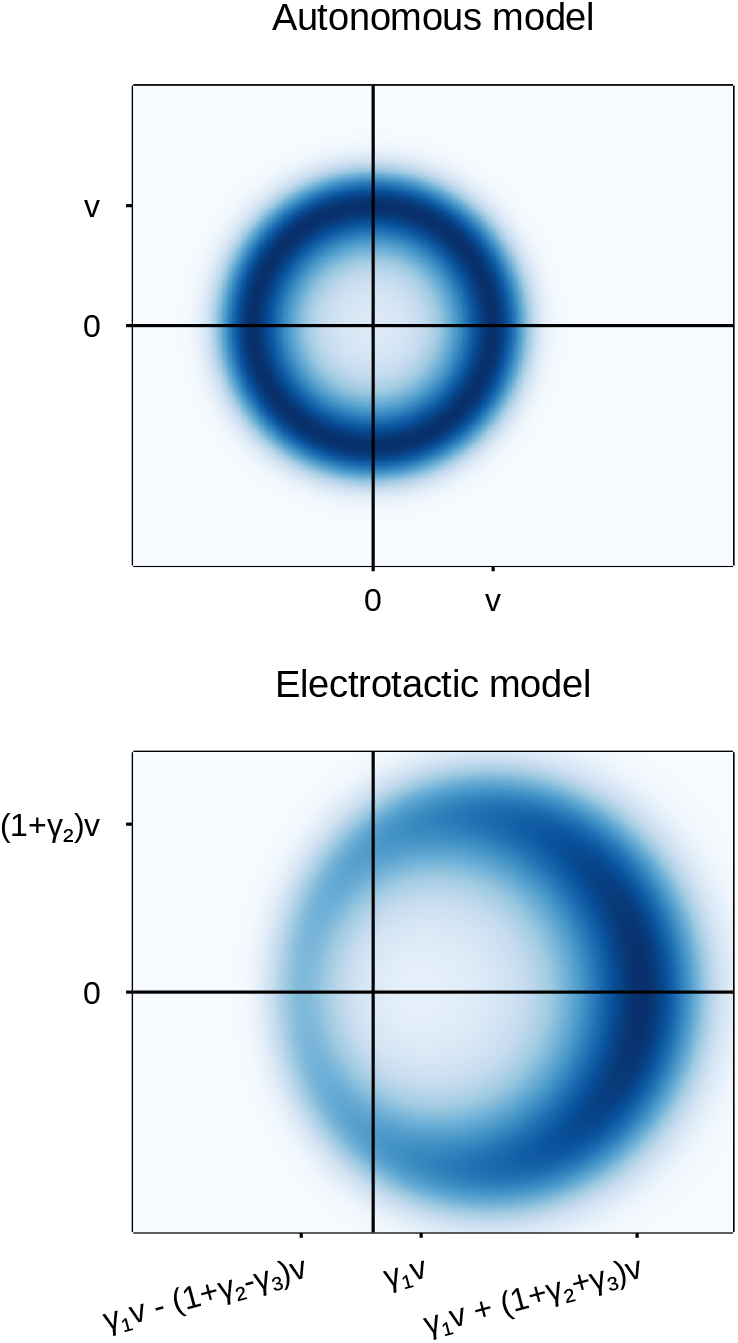
Comparison of the stationary distributions for the random velocity, **v**, under the autonomous and electrotactic models, where darker regions correspond to greater probability. The bottom plot shows the hypothesised electrotactic effects of an EF, applied in the positive *x* direction, parametrised by *γ*_1_,…, *γ*_4_. The effects of *γ*_1_, *γ*_2_ and *γ*_3_ are visible in the shape of the distribution. *Polarity bias* (*γ*_4_) produces asymmetry in the distribution density, shown as a darker region to the right of the figure.

### Outline

The primary goal of this work is to use single-cell experimental data to calibrate the parametrised mathematical model of spontaneous polarisation and electrotaxis. The model calibration process enables the identification of which of the four hypothesised electrotactic effects of EFs on cell motility can be observed in the experimental data. Importantly, the calibrated model also quantifies the relative contribution of each of these identified effects. The level at which we model the system allows us to subsequently use the calibrated model to simulate and predict the single-cell response to dynamic EFs.

The data used for model calibration is gathered from two assays in which the trajectories of motile human corneal epithelial cells are recorded for five hours: (a) without any EF applied for the entire experiment, and (b) with a DC EF at a reference strength of 200 mV mm^−1^, applied from left to right over hours 2–3 and from right to left over hours 4–5. These assays are termed the *autonomous* and *electrotactic* experiments, respectively. We use all five hours of the autonomous experiment and the first three hours of the electrotactic experiment as training data to calibrate the parameters of the autonomous and the electrotactic models. To calibrate the electrotactic model, we first identify which combination of the four hypothesised electrotactic effects is best supported by the data. After identifying which of the electrotactic effects are present in the model, we can then proceed to quantify the relative contribution of each of them to the observed electrotaxis induced by the EF.

After formulating and calibrating the extended model of electrotaxis, we use simulations of the calibrated model to predict how the cell trajectories evolve over the final two hours of the electrotactic experiment, in which the EF input has changed direction. We compare these predictions to the cell trajectories observed over the final two hours of the electrotactic experiment, held back to be used as test data, and thus validate the predictive capability of the model for dynamic EF inputs. The ability to make predictions of cellular motility using a calibrated, stochastic, uncertain model is a first step towards the future goal of model-based policy design for the electrotactic control of single-cell and population-level motility.

## MATERIALS AND METHODS

### Data collection

Two experiments were carried out, which we call the *autonomous* and *electrotactic* experiments. In both experiments, time-lapse images of human corneal epithelial cells, seeded at a low density, were acquired at 5 min intervals over 5 h. In the autonomous experiment, no EF was applied. In the electrotactic experiment, the cells were subjected to a DC EF at a reference strength, 200 mV mm^−1^, applied across the medium from *t* = 60 min to the end of the experiment. The EF was directed from left to right from 60 min to 180 min, at which point the field direction was reversed from right to left for 180 min to 300 min. Two replicates of each experiment were performed, with 26 and 27 cell centroids tracked for each of the autonomous assay replicates, and with 26 and 30 cell centroids tracked for each of the electrotactic assay replicates, all over the entire time horizon. Visual confirmation from the raw experimental output confirms that cell collisions were rare, due to the low density (100 cells cm^−2^) at which cells are initially seeded. We thus assume that cell–cell interactions can be neglected in the current model. The data used in this work is shared online at DOI:10.5281/zenodo.4749429.

We denote the resulting cell trajectory data **x**_noEF,*i*_(*t_k_*) and **x**_EF,*j*_(*t_k_*) for the autonomous and electrotactic experiments, respectively, where each trajectory is translated to begin at the origin, such that **x**_noEF,*i*_(0) = **x**_EF,*j*_(0) = 0 for all *i* and *j*. For these experiments, the indices *i* = 1,…, 53 and *j* = 1,…, 56 refer to the cell being traced, while *t_k_* = 5*k* min refers to the snapshot time points for *k* = 0,…, 60. We hold back **x**_EF,*j*_(*t_k_*) for *j* = 1,…, 56 and *k* = 36,…, 60 as test data, for the purposes of validating model predictions. The remaining data is used as training data, from which the model is calibrated. Thus, the training data consists of the trajectories from the autonomous experiment over the entire time horizon, and the trajectories from the electrotactic experiment over 0 min to 180 min, which are denoted by **x**_NoEF_ and **x**_EF_, respectively. The test data, denoted **x**_test_, consists of all trajectories from the electrotactic experiment over 180 min to 300 min, where the input EF has switched direction.

#### Materials

EpiLife culture medium with Ca^2+^ (60 μM), EpiLife defined growth supplement, and penicillin/streptomycin were purchased from ThermoFisher Scientific (Waltham, MA, USA). FNC Coating Mix was purchased from Athena Enzyme Systems (Baltimore, MD, USA). Dow Corning high-vacuum grease was purchased from ThermoFisher. Agar was purchased from MilliporeSigma (Burlington, MA, USA). Silver wires with 99.999% purity were purchased from Advent Research Materials Ltd. (Oxford, United Kingdom).

#### Cell culture

Telomerase-immortalized human corneal epithelial cells (hTCEpi) were routinely cultured in EpiLife medium supplemented with EpiLife defined growth supplement and 1% (v/v) penicillin/streptomycin. Cells were incubated at 37 °C with 5% CO_2_ until they reached ~70% confluence and were used between passages 55 and 65 for all cell migration assays.

#### Electrotaxis assay

Electrotaxis experiments were performed as previously described (15, 16) with minor changes. Briefly, the electrotaxis chambers (20 mm × 10 mm × 0.2 mm) were constructed in 100 mm petri dishes with glass strips and high-vacuum grease. The dimensions of the chambers were defined by the thickness and length of the glass slides, respectively. Chambers were coated with FNC Coating Mix, following the manufacturer’s instructions to facilitate cell attachment. Cells were seeded at a low density (100 cells cm^−2^) and cultured overnight (12 h to 18 h) in the chambers to allow sufficient attachment. Chambers were covered with glass coverslips and sealed with high-vacuum grease. Electric currents were applied to the chamber through agar-salt bridges connecting with silver–silver chloride electrodes in Steinberg’s solution (58 mM NaCl, 0.67 mM KCl and 0.44 mM Ca(NO_3_)_2_, 1.3 mM MgSO_4_ and 4.6 mM Tris base, pH 7.4). Fresh cell culture medium (Epilife) was added into reservoirs to ensure good salt bridge contact and to support cell viability during electric stimulation. An EF strength of 200 mV mm^−1^ was used unless otherwise noted. A pair of measuring electrodes was placed at the end of the electrotaxis chamber and connected to the multimeter for real-time monitoring of EF strength. The applied voltages were confirmed at the beginning of the experiment and every 30 min afterwards to ensure consistent EF application.

#### Time-lapse imaging and quantification of cell migration

Cell migration was monitored and recorded by phase-contrast microscopy using an inverted microscope (Carl Zeiss, Oberkochen, Germany) equipped with a motorized stage and a regular 10× objective lens. Time-lapse images were acquired at 5 min intervals using Metamorph NX imaging software (Molecular Device, Sunnyvale, CA, USA). To maintain standard cell culture conditions (37 °C, 5% CO_2_), a Carl Zeiss incubation system was used. Time-lapse images of cell migration were analyzed by using ImageJ software from the National Institutes of Health (http://rsbweb.nih.gov/ij/). Adherent cells in the images were manually tracked, and cells that divided, moved in and out of the field, or merged with other cells during the experiment were excluded from analysis. The position of a cell was defined by its centroid.

### Model construction

We constructed a mathematical model of single-cell dynamics. The model tracks the position of the cell centre in the plane, 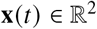, as a function of time, *t* ≥ 0 min, with initial condition **x**(0) = 0 at the origin. The position is a deterministic integral of cell velocity, **v**, such that

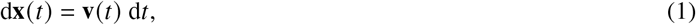

and the stochastic dynamics of **v** are modelled. The key to this modelling task is the non-dimensional internal variable representing the cell polarity, 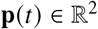. We assume that the polarity imparts a force on the cell that corresponds to its active motility, resulting in a velocity component **v**_cell_(*t*).

#### Modelling spontaneous polarisation and motility

We first describe the model of cellular motility with no biasing EF, which we will term the *autonomous* model. The only velocity component is that due to polarisation, so that we write the cell velocity as a single component,

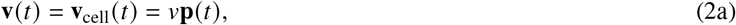

where the parameter *v* ≥ 0, with dimensions μm min^−1^, represents the modal magnitude of **v**_cell_ for a polarised cell. Note that Eq. (2a) implies that the polarity variable, **p**, is a non-dimensionalisation of the velocity component **v**_cell_. We further assume that the polarity, **p**, undergoes a random walk according to a Langevin diffusion, such that

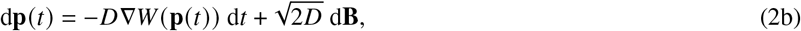

where 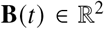 is a two-dimensional Wiener process, and the parameter *D* (in min^−1^) quantifies the speed at which the random walk approaches stationarity. The initial polarity, denoted **p**_0_ = **p**(0), also needs to be specified.

The *potential function W* (**p**) in Eq. (2b) is defined to capture the intended features of the autonomous model, namely that the magnitude of the cell velocity is randomly distributed around a modal value of *v*, and that the direction of the polarity is uniformly distributed at stationarity. It can be shown (17, 18) that the variability of the velocity around its modal value of *v* is determined by a non-dimensional *energy barrier*, denoted Δ*W*, that is sufficient to define the potential function, *W*(**p**). For further details on the definition of W, see the *Supplementary Material*. We will calibrate the autonomous model in Eq. (2) by identifying the parameters *v*, *D*, and Δ*W*.

#### Modelling motility bias due to an EF

We use a vector-valued function, **u**(*t*), with non-dimensional magnitude ||**u**(*t*)|| = *u*(*t*) to describe a (time-varying) DC EF of strength 200*u*(*t*) mV mm^−1^, directed parallel to **u**(*t*). In particular, the specific EF used in the electrotactic experiment, with magnitude 200 mV mm^−1^ in the positive *x* direction (left to right) over 60 min to 180 min, and reversed over 180 min to 300 min, is represented using the constant canonical unit vector, **i**, by the vector-valued function

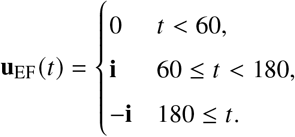

Note that the function **u**_EF_(*t*) represents the specific EF corresponding to the electrotactic experiment, while arbitrary EF inputs are modelled using the notation **u**(*t*). The autonomous model in Eq. (2) can be extended to include the four hypothesised effects of the EF. The *velocity bias* effect is accounted for by modelling the velocity using two components,

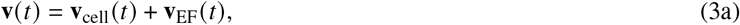

where the EF induces a deterministic velocity component in the direction of the field,

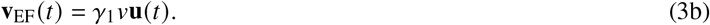

The two hypothesised electrotactic effects of *speed increase* and *speed alignment* are both modelled through adapting the velocity component induced by the cell polarity, originally defined in Eq. (2a), into

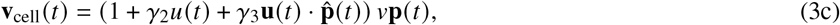

where 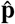 is the unit vector in the direction of the polarity, **p**. Finally, the hypothetical *polarity bias* effect is modelled in the stochastic evolution of the polarity variable **p**. We add a drift term proportional to the EF to the Langevin diffusion equation, such that

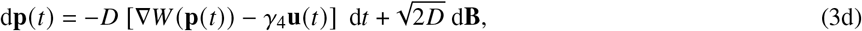

where *W*(**p**) is the same potential function as used in Eq. (2b). As for the autonomous model, the initial value for the polarity, denoted **p**_0_ = **p**(0), is also required.

Note that substituting **u**(*t*) ≡ 0 or setting *γ_i_* = 0 for all *i* = 1, 2, 3, 4 into Eq. (3) recovers the dynamics of the autonomous model, in Eq. (2). We will term the extended model in Eq. (3) the *electrotactic* model. It is parametrised by the three parameters *v*, Δ*W* and *D*, with the same meaning and dimensions as in the autonomous model, and also by *γ_i_* for *i* = 1, 2, 3, 4, which, since **u** is non-dimensional, are all non-dimensional.

The models in Eq. (2) and Eq. (3) result in a stochastic path for the velocity, **v**(*t*), with a parametrically determined stationary distribution. Following Eq. (1), each path can be integrated to produce a stochastic trajectory of the cell position over time. The stationary distributions of **v** under the autonomous and electrotactic models are depicted in Figure 1. The effect of each of the parameters *γ_i_*, and hence each of the hypothesised electrotactic effects, can be identified by comparing the position, scale, and asymmetries of the two stationary distributions.

#### Summarising simulations

For any given set of parameter values, *θ* = (*v*, Δ*W*, *D*, *γ*_1_, *γ*_2_, *γ*_3_, *γ*_4_), together with initial polarity, **p**_0_, and non-zero EF input, **u**(*t*), the stochastic model in Eq. (3) can be simulated. Note that, if the EF is zero, we simulate the autonomous model in Eq. (2). Each simulation produces a random trajectory, denoted *ω* = (**p**(*t*), **x**(*t*))_*t*≥0_. We will use *summary statistics* to analyse the model outputs by mapping each simulated trajectory, *ω*, to a number (or small set of numbers) that summarise the trajectory. More details of the summary statistics can be found in the *Supplementary Material*.

We define a set of summary statistics based on simulated cell positions at five-minute timepoints *t_j_* = 5 *j* over any given time interval, *t_n_* < *t_n+m_*. We consider: (a) the net horizontal cell displacement over the entire interval, (**x**(*t_n+m_*) − **x**(*t_n_*)) · **i**, denoted by *Y*_1_(*ω*); (b) the net absolute cell displacement over the entire interval, ||**x**(*t_n+m_*) − **x**(*t_n_*)||, denoted *Y*_2_(*ω*); (c) the path length, measured as the sum of displacements, 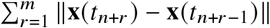, between the five-minute sample points, denoted *Y*_3_(*ω*), (d) the standard deviation of the displacements, ||**x**(*t_n+r_*) − **x**(*t*_*n*+*r*−1_)||, over *r* = 1,…, *m*, and denoted *Y*_4_(*ω*). Note that the four summary statistics *Y*_1_, *Y*_2_, *Y*_3_ and *Y*_4_ can also be applied to the observed data, **x**_NoEF,*i*_ and **x**_EF,*i*_, in addition to any simulated trajectory, *ω*.

In the models in Eq. (2) and Eq. (3), the polarity, **p**(*t*), evolves randomly from initial value **p**_0_. We define a further three summary statistics based on the simulated polarity, using a threshold polarity magnitude, 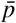. First, we define the *time to polarise*, *T*_1_, as the average time at which a simulated cell polarity, from initial polarity **p**_0_ = 0, first has polarity 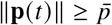. Conversely, we define the *time to depolarise*, *T*_0_, as the average time at which a simulated cell polarity, from initial polarity **p**_0_ = **i**, first has polarity 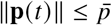. Finally, the value 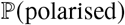 is defined as the probability that a simulated cell polarity at the end of an assay satisfies 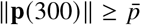. Note that these summary statistics *cannot* be applied to the observed data, since the polarity is not observed, and can only be used to summarise simulated trajectories.

### Model calibration and selection

Given the experimental training data sets, **x**_NoEF_ and **x**_EF_, the autonomous and electrotactic models can be *calibrated* by identifying the values of the parameters,

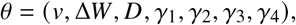

that are consistent with the observed behaviour. We employ a Bayesian approach to parameter inference, whereby prior beliefs about *θ*, encoded in a *prior distribution*, *π*(*θ*), are updated in the context of the experimental data according to Bayes’s rule,

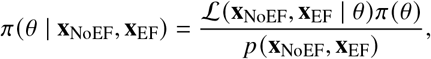

where 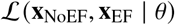 is the *likelihood* of observing the data under the models in Eq. (2) and Eq. (3) with the parameter value *θ*. The resulting *posterior distribution*, *π*(*θ* | **x**_NoEF_, **x**_EF_), represents the remaining uncertainty in the parameter values, given the training data (19).

The simulation and inference algorithms used in this work have been developed in Julia 1.5.1 (20). The code is publicly available at github.com/tpprescott/electro.

#### Bayesian synthetic likelihoods and sequential Monte Carlo

In practice, the likelihood cannot be calculated directly, and so we require a likelihood-free approach. We replace the true likelihood with a *synthetic likelihood*, where for each value of *θ* the likelihood is approximated by the likelihood of summarised data under an empirical Gaussian distribution. The empirical distribution is fit to the sample mean and covariance of a set of n *=* 500 summarised simulations (21–23). The summary statistics we use are *Y*_1_, *Y*_2_, *Y*_3_ and *Y*_4_, as described in *Summarising simulations*, above. We summarise the interval 0 min to 300 min for the trajectories from the autonomous experiment, and the separate intervals 0 min to 60 min and 60 min to 180 min for the trajectories from the electrotactic experiment. To mitigate the computational burden of the large number of model simulations required for parameter inference, we combine a sequential Monte Carlo (SMC) algorithm with synthetic likelihoods (21, 24, 25). This approach is a popular strategy for efficiently sampling from a target distribution, and also allows the exploitation of parallelisation to speed inference (21, 25–27). We provide full details of the SMC inference approach using summary statistics and synthetic likelihoods in the *Supplementary Material*.

#### Prior specification and model selection

The space of possible parameter values is defined as the product of intervals,

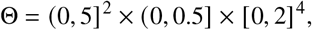

where the interval bounds were chosen based on a preliminary qualitative, visual analysis of the simulation outputs in comparison to observed data. In order to identify which of the sixteen possible combinations of the four hypothesised electrotactic effects are best supported by the experimental data, we will define sixteen possible priors on Θ. For each of the sixteen subsets, *X* ⊆ {1, 2, 3, 4}, we define a uniform prior distribution *π_x_* (*θ*) on Θ that takes a constant, positive value for parameter vectors *θ* if and only if *γ_i_* > 0 for all *i* ∈ *X*, and *γ_i_* = 0 otherwise. Thus, by performing Bayesian inference using the prior distribution, *π_x_*, we constrain the electrotactic model in Eq. (3) to model only electrotactic effects included in the subset *X* ⊆ {1, 2, 3, 4}.

We define an optimisation problem that aims to prevent over-fitting, by balancing the closeness of the model fit to data while prioritising smaller parameter dimensions. The optimal subset, *X*, of electrotactic effects is defined as the maximiser of the objective function,

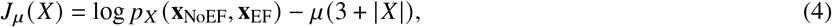

where the regularisation parameter *μ* > 0 controls the cost of over-fitting by penalising the total number of non-zero parameters. This number is three, corresponding to *v*, Δ*W*, and *D*, plus |*X*|, corresponding to the positive *γ_i_* for *i* ∈ *X*. We use *μ* = 0 and *μ* = 2 in our analysis, though we note that the choice of *μ* is somewhat arbitrary. Choosing *μ* = 2 imposes a penalty on the parameter dimension analogous to that used in the Akaike information criterion (19). One interpretation of the value of *μ* is that it effectively imposes a ‘prior’ on the subsets, *X* ⊆ {1, 2, 3, 4}, with probability mass proportional to exp(−*μ*|*X*|).

The first term in *J_μ_*(*X*) measures the closeness of fit between the data and the model, when constrained to only include the electrotactic effects in *X*. This fit is defined for each *X* ⊆ {1, 2, 3, 4} by the value of the partition function,

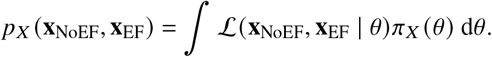

As the likelihoods 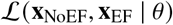 cannot be calculated directly, the partition functions *p_x_*(**x**_NoEF_, **x**_EF_) are estimated for each *X* by Monte Carlo sampling, where again the simulation-based synthetic likelihood is used in place of the true likelihood. More details of the specific sequential Monte Carlo sampling methodology used for this estimate are given in the *Supplementary Material*.

## RESULTS

We initially calibrate the autonomous model, based on the portion of the training data set **x**_NoEF_ from the autonomous experiment alone, in order to confirm the principle of the modelling framework and its ability to replicate observed behaviours, and to check that the parameters are identifiable from the data. Then, we calibrate the full electrotactic model using the full training data set, **x**_NoEF_ and **x**_EF_, in two stages. We first assess which subset of the four hypothesised electrotactic effects are best supported by the data. After choosing the optimal combination of electrotactic effects, we then calibrate the parameters of the selected electrotactic model.

### Parameters of the autonomous model are identifiable

We begin by confirming that the chosen modelling and inference approaches appropriately capture the autonomous experimental behaviour, **x**_NoEF_, where no external EF is applied. The cell trajectories in this portion of the training data are depicted in Figure 2(a). This scenario is modelled by the autonomous model in Eq. (2), which depends on three parameters, *θ*_NoEF_ = (*v*, Δ*W, D*). The Bayesian synthetic likelihood approach was used to generate posterior samples for: the characteristic speed of a polarised cell, *v* μmmin^−1^; the timescale constant, *D* min^−1^, which determines the characteristic timescale of the spontaneous polarisation dynamics; and the dimensionless parameter, Δ*W*, which determines the variability of the cell polarity around its modal value.

**Figure 2:**
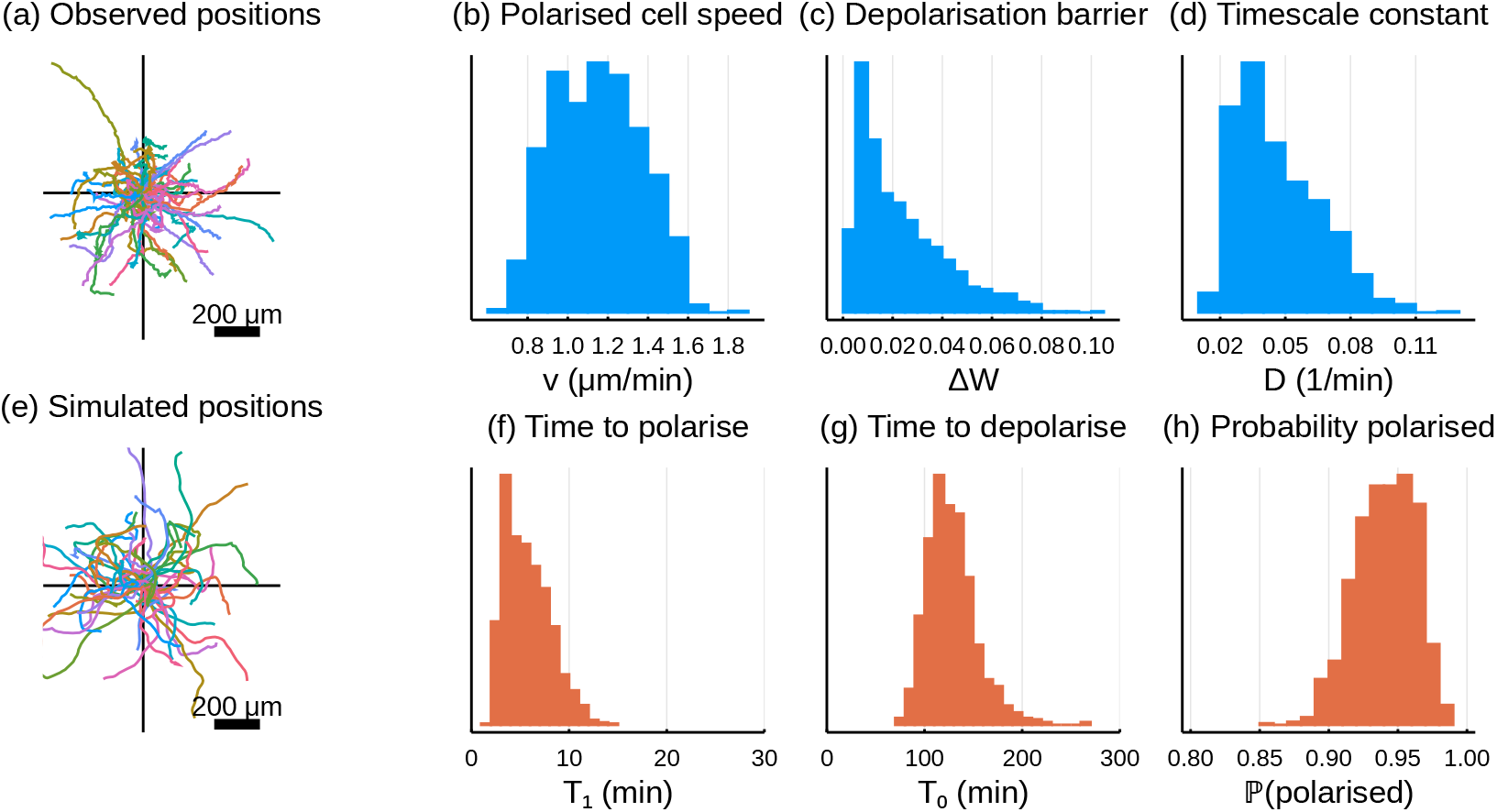
Parameter inference and simulation of the autonomous model. (a) The subset of the training data used to calibrate the autonomous model, corresponding to all observed trajectories under the autonomous experiment. (b–d) All one-dimensional projections of the posterior sample from *π* (*θ*_NoEF_ | **x**_NoEF_). The covariance structure of the posterior is given in *Supplementary Material*, Figure S2. (e) Simulations of the calibrated model, using parameters randomly selected from the posterior depicted in (b–d). (f–h) Posterior predictive samples for *T*_1_ (time to polarisation), *T*_0_ (time to depolarisation), and 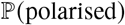 (probability of a cell being polarised at the final time) for simulations from the autonomous model with parameters taken from the posterior, *π*(*θ*_NoEF_ | **x**_NoEF_), depicted in (b–d).

Figure 2(b–d) depicts the marginals of the posterior distribution, *π*(*θ*_NoEF_ | **x**_NoEF_), for each of the three calibrated parameters. The prior distribution used for Bayesian inference assumed that the parameters were independently uniformly distributed on the intervals 0 < *v* ≤ 5 μmmin^−1^, 0 < Δ*W* ≤ 5, and 0 < *D* ≤ 0.5 min^−1^. Each plot in Figure 2(b–d) demonstrates that the posteriors are concentrated within a small interval of the prior support, implying that the parameters of the autonomous model are identifiable from the experimental data, with quantifiable uncertainty.

The sample median parameter value, calculated from the sample in Figure 2(b–d), can be used as a point estimate for the parameter values: *v* = 1.15 μmmin^−1^, Δ*W* = 0.018, and *D* = 0.042 min^−1^. In Figure 2(e), we depict a random sample of trajectories simulated from the autonomous model, with parameter values sampled from the empirical posterior, that compare well with the observed trajectories from the autonomous experiment, **x**_NoEF_. A comparison between these plots shows that parameter inference based only on the selected four-dimensional summary statistics produces a close match (for this point estimate) between the visual characteristics of simulations and experimental observations. A more detailed analysis of the fit of the calibrated autonomous model to the training data is given in the *Supplementary Material*, including a comparison of posteriors trained on the two replicates separately, and a cross-validation of the posterior predictive distribution of the four summary statistics.

Figure 2(b–d) quantifies the uncertainty in each parameter value resulting from the Bayesian approach to parameter inference. In order to make sense of this uncertainty in terms of the model outputs, simulations can be used to interpret how the uncertainty propagates to observable behaviour. Figure 2(f–h) depicts an estimate of the uncertainty in (f) the average time a simulated cell takes to polarise, *T*_1_, (g) the average time a simulated cell takes to depolarise, *T*_0_, and (h) the proportion of simulated cells that are polarised by the end of the experiment, 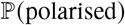. Each of these distributions are conditioned on the posterior parameter distribution in Figure 2(b–d). This procedure allows us to map quantified uncertainty in the parameter values to uncertainty in cell behaviour. The calibrated model suggests that the expected time for a cell to spontaneously polarise (i.e. without an EF applied) ranges from 2.8 min to 10 min (5% to 95% quantiles), with median value of 5.4 min. Similarly, the expected time for a cell to depolarise is 94 min to 174 min, with median value 124 min. Finally, the probability that a simulated cell is polarised (in any direction) at the end of the experiment is 0.9 to 0.97, with median value 0.94.

### One of the four proposed electrotactic effects is supported by the data

Given that the autonomous model can be calibrated to the data set from the autonomous experiment, we now seek to calibrate the full electrotactic model to the entire training data set taken from both experiments. However, some or all of the hypothesised electrotactic effects used to define the model in Eq. (3) may not be supported by the experimental data. Thus, we first use the training data to select which of these proposed effects can be detected in the observed cell behaviours. Recall that the parameters *γ*_1_, *γ*_2_, *γ*_3_ and *γ*_4_ correspond to four distinct hypothesised electrotactic effects: velocity bias, speed increase, speed alignment, and polarity bias. Positive values of the parameters *γ*_i_, for *i* = 1,2,3,4, mean that the corresponding effect is included in the model. Conversely, setting any of these parameters to zero excludes the corresponding effect(s) from the model. There are a total of 2^4^ = 16 possible combinations of the four proposed electrotactic effects that the model in Eq. (3) can implement, through combinations of positive and zero parameter values.

Each of the 16 possible combinations of the four electrotactic effects corresponds to a subset *X* ⊆ {1,2,3,4}. We evaluate each combination of electrotactic effects, given by *X*, with respect to the objective function, *J_μ_*(*X*), given in Eq. (4). This objective quantifies the trade-off between the model fit and the number of non-zero parameters in order to select a suitably accurate model while avoiding over-parametrisation. Figure 3 ranks each of the 16 possible combinations of electrotactic effects, *X* ⊆ {1,2,3,4}, using two different objective functions. The top plot considers *μ* = 0, such that the maximiser of *J*_0_ is the combination that gives the best fit to data, with no consideration given to the dimension of parameter space. The bottom plot uses *μ* = 2, which imposes a marginal cost on increasing the dimension of parameter space. Both objective functions are maximised by the singleton subset *X* = {4}, by a margin of over 10 from *X* = {1,4} in second-place. This margin implies a Bayes factor (19) greater than 10 in favour of a prior such that *γ*_4_ ∈ (0,2], with *γ*_1_ = *γ*_2_ = *γ*_3_ = 0, thus providing strong support for including only the polarity bias effect of the EF in our model, and neglecting all of the other hypothesised effects. Indeed, we can also conversely conclude from Figure 3 that any prior that sets *γ*_4_ = 0 would induce a poor fit to the observed data.

**Figure 3:**
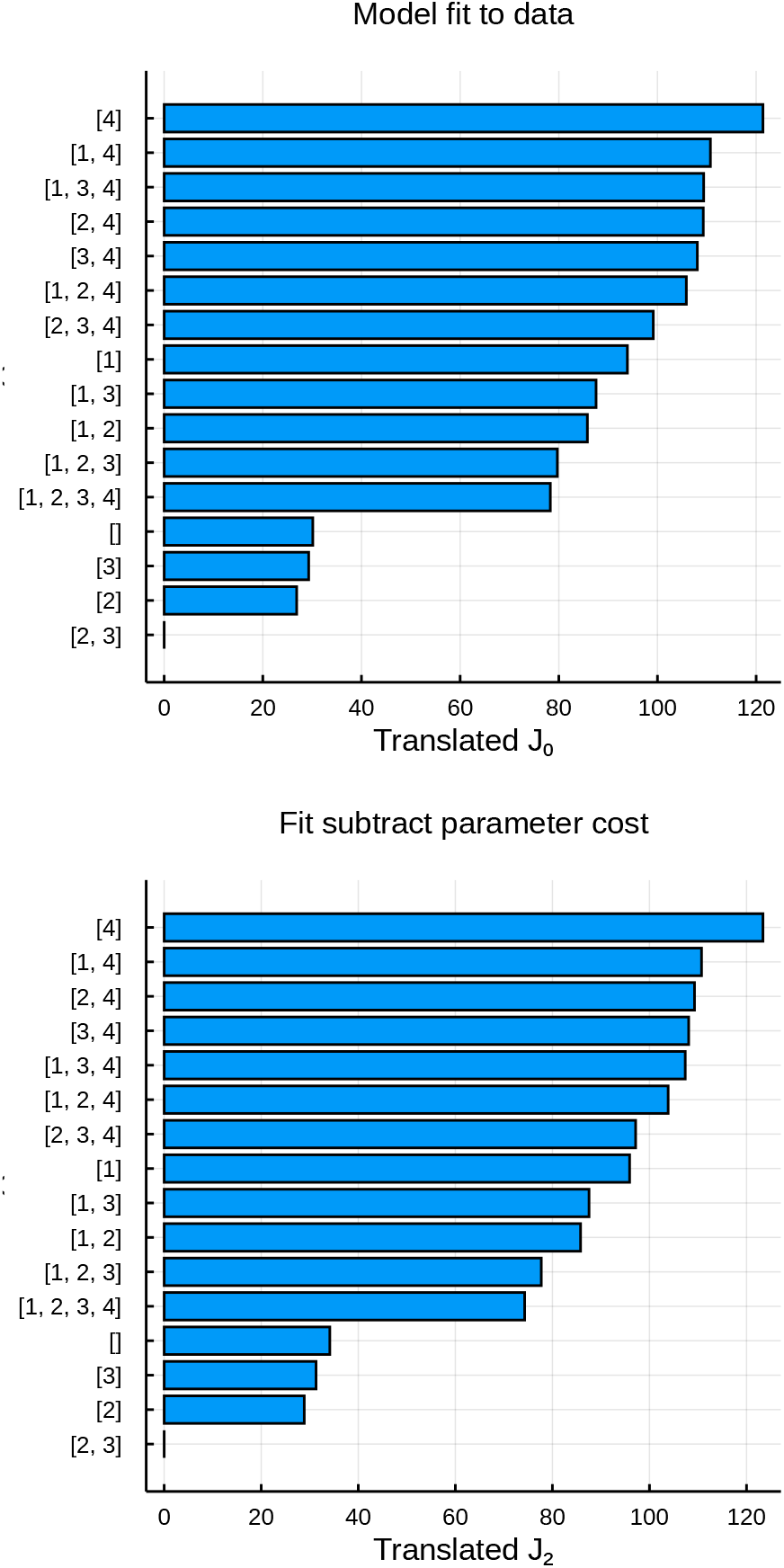
Objective functions *J*_0_ (*X*) and *J*_2_ (*X*) from Eq. (4), for combinations of electrotactic effects indexed by *X* ⊆ {1,2,3,4}. Greater values are preferred. Each objective function is translated to have zero minimum value.

### The electrotactic effects of the EF on motility can be quantified

Recall that cell motility is modelled as the sum of an active force component, deriving from cell polarisation, and a component comprised of other external forces acting on the cell. In the selected model found in the preceding section, we found *γ*_1_ = 0, meaning that the training data provides no evidence that the EF imparts an external force. Finding that *γ*_2_ = *γ*_3_ = 0 further implies that polarised cells do not travel any faster in the presence of a field, neither uniformly nor only if polarised in alignment with the field. Instead, the EF produces the observed bias in cell motility solely due to causing cells to preferentially polarise in the direction of the EF. In this section, we calibrate the electrotactic model by using the entire training data set, **x**_NoEF_ and **x**_EF_, to infer the posterior distribution of *γ*_4_ > 0, while also refining the posterior distributions of *v*, Δ*W*, and *D*.

Bayesian synthetic likelihoods were used to calibrate the electrotactic model by inferring the posterior distribution, *π*(*θ* | **x**_NoEF_, **x**_EF_), for

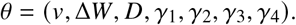

The chosen prior distribution, *π*_{4}_(*θ*), is the product of independent uniform distributions on the intervals 0 < *v* ≤ 5 μm min^−1^, 0 < Δ*W* ≤ 5, and 0 < *D* ≤ 0.5 min^−1^, multiplied by an independent and uniformly distributed prior for the polarity bias parameter on the interval 0 < *γ*_4_ ≤ 2. The remaining parameters in *π*_**x**_(*θ*) are fixed at *γ*_1_ = *γ*_2_ = *γ*_3_ = 0.

Figure 4(a–d) shows the empirical marginals from the posterior sample from *π*(*θ*| **x**_NoEF_, **x**_EF_), constructed using Bayesian synthetic likelihoods and SMC sampling. The marginals shown correspond to the four non-zero parameters of the model, *v*, Δ*W, D*, and *γ*_4_. The posterior marginal distributions of the previously inferred parameters, *v*, Δ*W* and *D*, closely match those in Figure 2(b–d), as depicted in Figure S7 of the *Supplementary Material.* Similarly to Figure 2, the posterior distribution is concentrated in a small region of the prior domain, providing evidence that each of the parameters is identifiable using the chosen summary statistics.

**Figure 4:**
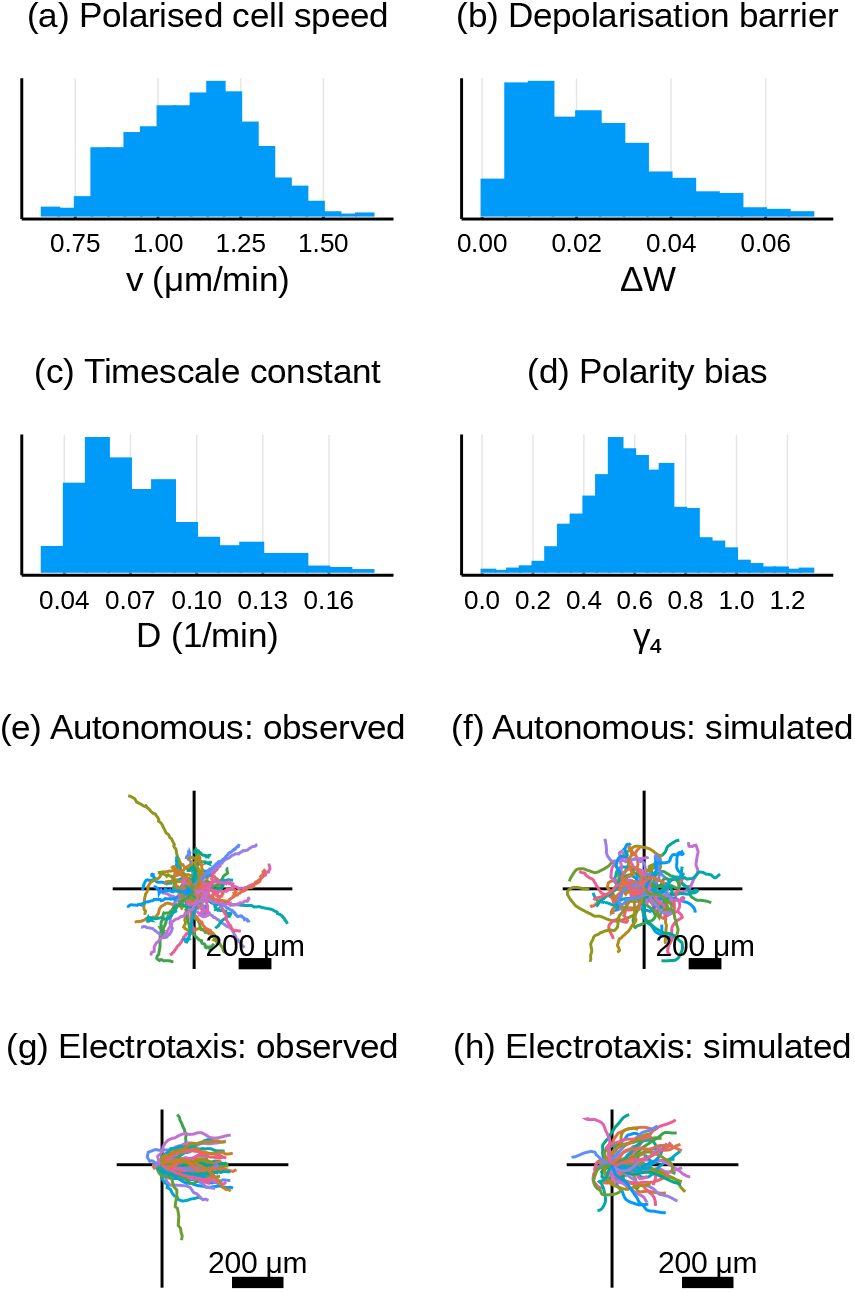
Empirical posterior samples inferred from training data: all trajectories from the autonomous experiment, and also from the electrotactic experiment for 0 min to 180 min. (a–d) One-dimensional projections of the empirical posterior distribution for all non-zero parameter values, based on the selected prior, *π*_[4]_. In the *Supplementary Material*, the two-dimensional projections of this posterior are depicted in Figure S6, and a comparison of the posterior distributions of *v*, Δ*W* and *D* in Figure 2(b–d) and here is depicted in Figure S7. (e–h) Observed and simulated trajectories for (e, f) the autonomous experiment, and (g, h) the electrotactic experiment over 0 min to 180 min. Simulations in (f,h) were produced for randomly sampled parameter values from the posterior in (a–d).

The median of the SMC sample depicted in Figure 4(a–d) can be used as a point estimate for the parameter values: *v* = 1.11 μmmin^−1^, Δ*W* = 0.020, *D* = 0.069 min^−1^, *γ*_4_ = 0.60. Figure 4(e–h) compares the training data against 50 simulations from both models, Eq. (2) and Eq. (3), using parameter values randomly sampled from the empirical posterior depicted in Figure 4(a–d). The observed bias in motility towards the direction of the EF is reflected in the stochastically simulated outputs. This provides visual confirmation that parameters inferred by Bayesian synthetic likelihood, based on the chosen summary statistics, produce simulated outputs that share observable characteristics with the experimental data. In the *Supplementary Material* we provide a more detailed validation of the fit of the calibrated model to the training data, including a cross-validation of the posterior predictive distributions of the summary statistics against the observed summary statistics, in Figure S8.

### Validation against test data

Recall that the portion of the data collected from the electrotactic experiment corresponding to the time interval 180 min to 300 min was held back from the training set used to calibrate the model. The predictions of the calibrated electrotactic model can be validated against this test data set, as a prediction of the cell responses to a reversed EF input.

In Figure 5, we compare the predictions to the test data in two ways. Figure 5 depicts the marginals of the empirical posterior predictive distribution for all four summary statistics over the interval 180 min to 300 min, overlaid against the corresponding summarised test data. These distributions show a good level of agreement, demonstrating that these characteristics of the observed motility data can be predicted by the model. In Figure 5(e–f) we also compare the observed trajectories over 180 min to 300 min (translated to begin at the origin) against a set of simulated trajectories, using parameter values sampled randomly from the posterior in Figure 4(a–d). A visual comparison shows that the diversity of observed trajectory characteristics is well-captured by the stochastic simulations and parameter uncertainty. These comparisons provide evidence helping to validate the calibrated model, by demonstrating its ability to accurately predict the cellular response to dynamic EF inputs against unseen data.

**Figure 5:**
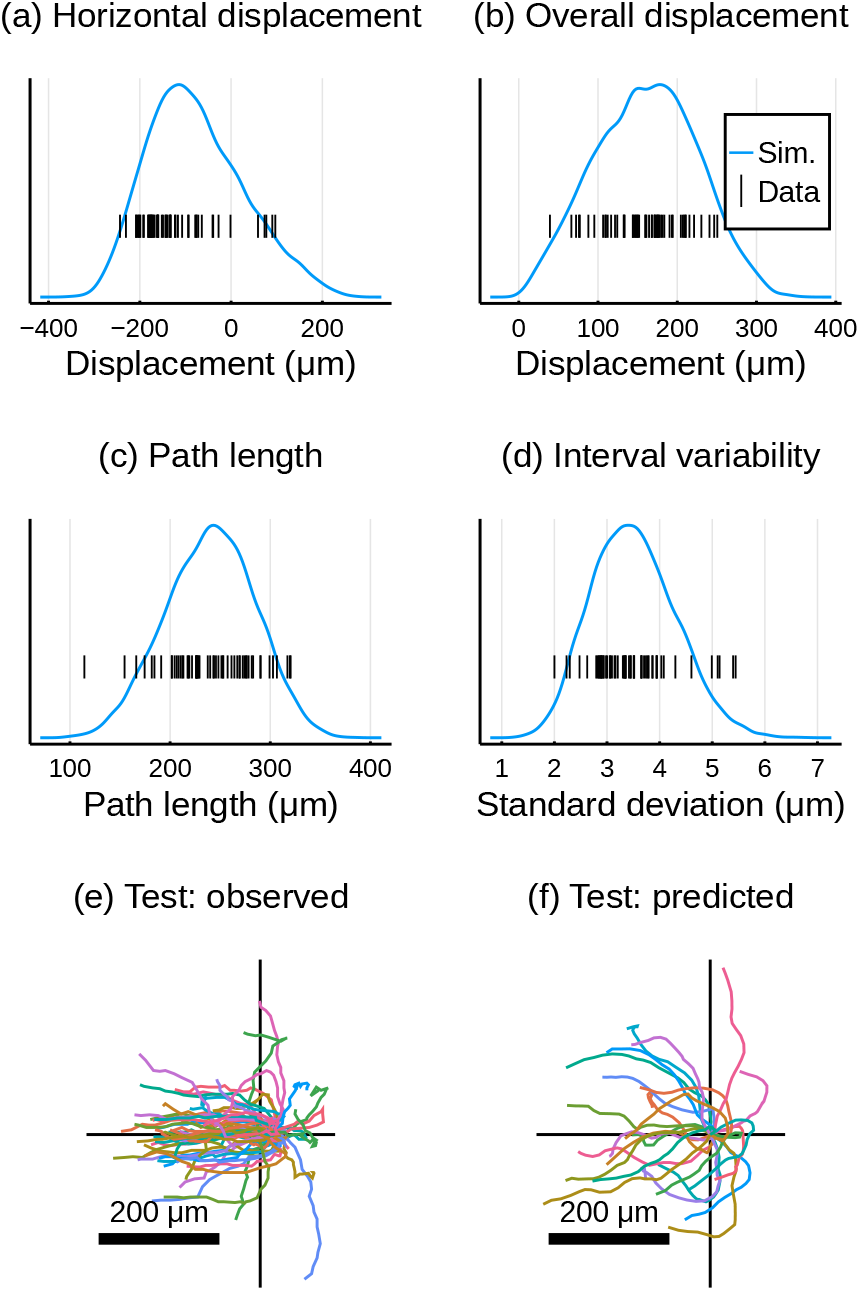
Comparing predictions of the calibrated electrotactic model against test data: observed trajectories from the electrotactic experiment for 180 min to 300 min. (a–d) One-dimensional projections of the summary statistics. Curves are an empirical distribution of simulated summary statistics for each parameter value from the posterior sample depicted in Figure 4(a–d). We overlay a barcode plot of each of the summary statistics of the observed test data. (e–f) Observed and simulated trajectories over 180 min to 300 min. Simulations in (f) were produced for randomly sampled parameter values from the posterior in Figure 4(a–d).

## DISCUSSION & CONCLUSION

The primary goal of this work has been to use mathematical modelling to quantitatively identify the contributions of multiple hypothesised means by which EFs induce electrotaxis in single cells. We have presented an empirical, parametrised, agent-based model of electrotactic cell motility, and shown that it can be calibrated to single-cell trajectory data using likelihood-free Bayesian inference. To our knowledge, although many models of single-cell and collective motility under environmental cues have been developed (13), there have been few mathematical models of electrotaxis (28, 29), and this work is the first use of detailed mathematical modelling at a single-cell level to quantify motility under electrotaxis. Moreover, the inferred parameter values of the calibrated model provide quantitative, mechanistic insights into experimentally-observed electrotaxis.

Specifically, by calibrating the model to experimental observations of electrotaxis in human corneal epithelial cells, we have concluded that the observed bias in motility is the result of *polarity bias*, where cells preferentially evolve the direction of their polarisation to align with the direction of the EF. The data does not support the hypothesis that an EF contributes an external force on the cell; nor that polarised cells travel at different speeds in the presence of a field. By carefully calibrating the parametrised mathematical model to experimental data, we have quantified the effect of the polarity bias on the electrotactic phenotype of this cell line.

A key strength of the model presented in Eq. (3) is its flexibility. The parametric design means that the Bayesian calibration methodology used in this work can be recapitulated to calibrate the same model to electrotaxis assays using other cell types or with different experimental conditions. Thus, observed differences in spontaneous and electrotactic motility between different cells and experimental conditions (3, 5) can be modelled and predicted within a common parametric framework. It is also important to acknowledge that we have chosen from only four hypothetical observable effects of electrotaxis. Other electrotactic effects may be reasonably included in the modelling process: for example, the EF may induce changes to the rate of polarisation and depolarisation (3). The electrotaxis model can straightforwardly be extended and recalibrated to account for any alternative hypothetical effects.

We have also considered EFs at a single reference strength, requiring a single parameter to quantify each hypothesised electrotactic effect. However, the characteristics of electrotaxis have been observed to vary nonlinearly with EF strength (5). The model is sufficiently flexible to account for this phenomenon through the replacement of the parameters *γ_i_* with functions Γ_*i*_ (*u*) that vary with the EF strength, *u* mV mm^−1^. The challenge will then be to use experimental data gathered from assays using EFs of different strengths to infer each of the functions Γ_*i*_ in place of each of the parameters *γ_i_*.

The model we have presented predicts single-cell electrotactic behaviour. However, there is a wealth of data and analysis on electrotaxis in the context of cell populations (3, 4, 6–9, 13). The electrotaxis model in this paper is a starting point for a comprehensive agent-based model that also incorporates phenomena such as volume exclusion, adhesion, elastic collisions, contact inhibition, and so on (13, 30, 31). Furthermore, there is significant scope for linking the calibrated parameters of the single-cell model described in this work to the construction of lower-level models of the intracellular processes that give rise to electrotaxis. Multifidelity approaches (27, 32) that can link experiments and information at the intracellular, single-cell and multicellular level will be vital to identify and quantify the biasing effects of EFs on the collective motility of cell populations (12, 14, 33).

The model considered in this paper, and the Bayesian uncertainty quantification of its parameters, are important tools for enabling stochastic model predictive control designs of such policies based on output feedback and filtering (34). We have therefore provided a significant step towards the real-time model predictive control of populations of electrotactic cells.

## AUTHOR CONTRIBUTIONS

The experiments and image analysis were performed by KZ and MZ. The modelling, simulation and inference were carried out by TPP and REB. All authors contributed to writing the manuscript.

## ACKNOWLEDGMENTS

The experiments in the Zhao lab were supported by the AFOSR-MURI grant FA9550-16-1-0052 and NIH grant 1R01EY019101. TPP and REB are supported by BBSRC through grant BB/R000816/1, and REB is supported by a Royal Society Wolfson Research Merit Award.

## Notes

https://github.com/tpprescott/electro

https://zenodo.org/record/4749429

### Competing Interest Statement

The authors have declared no competing interest.

### Summary of Updates

Additional data collected; conclusions on model calibration and parameter values updated in the light of new data; model predictions validated against held-back test sets.

