## Supplementary Text and Figures for "Quantifying the impact of electric fields on single-cell motility"

TP Prescott K Zhu M Zhao RE Baker

### 1 Preliminary data analysis

To justify the assumption that the cell velocity evolves according to a random walk, we performed an initial analysis of both control data replicates. In Figure S1(a,b) we plot the distribution of displacement distances travelled in each five-minute sampling interval over the five-hour experiment, smoothed using a kernel density estimate, and where each curve corresponds to a different cell. These distributions in displacement distances are consistent with the modelling assumption that cells can spontaneously polarise, in which state they travel at a positive speed, and can also transiently depolarise.

Furthermore, we can also justify our assumption that the direction of motion of each cell drifts stochastically over time. Figure S1(c,d) plots the autocorrelations between the observed angle of cell displacements over each five-minute interval for each cell trajectory, measured for time lags from 5 min to 60 min. This figure demonstrates that the autocorrelation in the observed motility direction decays with increasing time lag. This observation is consistent with the modelling assumption that the velocity direction evolves according to a random walk.

### 2 Mathematical model of electrotaxis

The autonomous model of cellular velocity in Eq. (2) is given by

$$\mathbf{v}(t) = v\mathbf{p}(t), \tag{2a}$$

$$\mathrm{d}\mathbf{p}(t) = -D\nabla W(\mathbf{p}(t))\ \mathrm{d}t + \sqrt{2D}\ \mathrm{d}\mathbf{B}, \tag{2b}$$

and the electrotactic model in Eq. (3) is given by

$$\mathbf{v}(t) = \mathbf{v}_{\mathrm{cell}} + \mathbf{v}_{\mathrm{EF}}, \tag{3a}$$

$$\mathbf{v}_{\mathrm{EF}}(t) = \gamma_1 v \mathbf{u}(t), \tag{3b}$$

$$\mathbf{v}_{\mathrm{cell}}(t) = (1 + \gamma_2 u(t) + \gamma_3 \mathbf{u}(t) \cdot \hat{\mathbf{p}}(t))\, v\mathbf{p}(t), \tag{3c}$$

$$\mathrm{d}\mathbf{p}(t) = -D\left(\nabla W(\mathbf{p}(t)) - \gamma_4 \mathbf{u}(t)\right)\ \mathrm{d}t + \sqrt{2D}\ \mathrm{d}\mathbf{B}. \tag{3d}$$

Here, cell velocity is denoted by $\mathbf{v}$ and cell polarity is denoted by $\mathbf{p}$. The vector $\mathbf{u}$ represents the EF with magnitude $\|\mathbf{u}(t)\| = u(t)$, scaled such that $u(t) = \alpha$ represents a field of strength $200\alpha$ $\mathrm{mV\,mm^{-1}}$ applied at time $t$ in the direction parallel with $\mathbf{u}(t)$. The two-dimensional standard Wiener process is denoted by $\mathbf{B}$, and $\hat{\mathbf{p}}$ is the unit vector in the direction of polarity. Both models depend on the parameters $v$, with units $\mu\mathrm{m\,min^{-1}}$, and $D$, with units $\mathrm{min^{-1}}$. Thus $\mathbf{p}$ is a non-dimensional quantity. The additional parameters, $\gamma_1, \ldots, \gamma_4$, in the electrotactic model parametrise the four hypothesised electrotactic effects, as described in the main text.

Also common to both models is the potential function $W(\mathbf{p})$. This function is defined to capture the intended features of the autonomous model, namely that cells stochastically and spontaneously polarise, and that the direction of the polarity is uniformly distributed at stationarity. Denoting $p = |\mathbf{p}|$, it follows from the latter requirement that the potential function $W(\mathbf{p}) = W(p)$ must be radially symmetric. The interpretation of the parameter $v$ as the modal speed of a polarised cell also implies that the polarised state is characterised by $\mathbf{p}$ stochastically evolving in the regime $p \approx 1$. We therefore require a potential function with local minimum at $p = 1$. Following [11], this function is implemented as

$$W(p) = \beta \left( \frac{1}{4} p^4 - \frac{1}{2} p^2 \right), \tag{S1}$$

where $\beta > 0$ defines the local minimum valueof the wellat $p = 1$. It can be shown [17, 18] that the rate at which the cell transiently depolarises is determined solely by the timescale parameter, $D$ min$^{-1}$, and the non-dimensional value of the *energy barrier*, $\Delta W = \beta/4$. Hence, we calibrate the models in Eq. (2) and Eq. (3) by inferring the common parameters, $v$, $D$, and $\Delta W$, together with the parameters $\gamma_1, \dots, \gamma_4$ specific to the electrotactic model.

**Note on polarity definition** Our description of the model interprets the variable $\mathbf{p}$ as the cell polarity, and treats velocity as the combination of a polarity component and a component due to the EF. Another interpretation of $\mathbf{p}$ is available if we specifically define single-cell polarity as the non-dimensionalisation of the velocity by $v$. In the electrotactic model, this alternative definition identifies cell polarity as the variable

$$\mathbf{v}/v = (1 + \gamma_2 u + \gamma_3 \mathbf{u} \cdot \hat{\mathbf{p}})\mathbf{p} + \gamma_1 \mathbf{u}.$$

The variable $\mathbf{p}$, with dynamics (3d), is then interpreted as a slowly-responding component of the cell polarity (in the alternative definition) to the EF input, while $\gamma_1 \mathbf{u}$, identifiable with *velocity bias*, is an instantly-responding component of the cell polarity. However, these definitions are internal to the model, in the sense that they have no effect on the observable position or velocity of simulated cells. Thus, in the current work, we choose to identify 'cell polarity' as the modelled variable $\mathbf{p}$, while noting that alternative interpretations are possible.

### 3 Likelihood-free Bayesian inference

To infer the parameters of the model, we will use the data from both experimental assays. We use all recorded cell positions from the autonomous experiment over $t \in [0, 300]$, denoting this data set by $\mathbf{x}_{\text{NoEF}}$. In addition, we use the recorded cell positions from the electrotactic experiment, but only over $t \leq 180$ min, denoting this data set by $\mathbf{x}_{\text{EF}}$. Note that the data from the electrotactic experiment gathered over $t > 180$ min will be held back as test data. The Bayesian inference framework uses the experimental training data, $\mathbf{x}_{\text{NoEF}}$ and $\mathbf{x}_{\text{EF}}$, to update a prior distribution, $\pi(\theta)$, into a posterior distribution, $\pi(\theta \mid \mathbf{x}_{\text{NoEF}}, \mathbf{x}_{\text{EF}})$, by multiplying by the *likelihood*, $\mathcal{L}(\mathbf{x}_{\text{NoEF}}, \mathbf{x}_{\text{EF}} \mid \theta)$, according to Bayes' rule,

$$\pi(\theta \mid \mathbf{x}_{\text{NoEF}}, \mathbf{x}_{\text{EF}}) \propto \mathcal{L}(\mathbf{x}_{\text{NoEF}}, \mathbf{x}_{\text{EF}} \mid \theta) \pi(\theta).$$

To define the likelihood, we first consider simulations of the models in Eq. (2) and Eq. (3).

For a given parameter vector, $\theta$, initial polarity, $\mathbf{p}_0$, and non-zero EF input, $\mathbf{u}(t)$, the model in Eq. (3) is simulated and a trajectory, $\omega = (\mathbf{p}(t), \mathbf{x}(t))_{t \geq 0}$, is produced. This stochastic trajectory has conditional density $p(\omega \mid \theta, \mathbf{p}_0, \mathbf{u}(t))$. We assume that there is a known distribution, $\varphi(\mathbf{p}_0)$, for the initial polarity: for the inference procedure carried out in the main text, we assume that $\varphi$ is a Gaussian distribution with zero mean and diagonal covariance matrix, with component-wise variances of 0.1. For the two specific experimental inputs, $\mathbf{u}_{\text{NoEF}}(t) \equiv 0$

and

$$\mathbf{u}_{\mathrm{EF}}(t) = \begin{cases} 0 & t < 60, \\ \mathbf{i} & 60 \leq t < 180, \\ -\mathbf{i} & 180 \leq t, \end{cases}$$

we integrate the density $p$ with respect to $\varphi(\mathbf{p}_0)$ and thus define two densities,

$$p_{\mathrm{NoEF}}(\omega \mid \theta) = \int p(\omega \mid \theta, \mathbf{p}_0, \mathbf{u}_{\mathrm{NoEF}})\varphi(\mathbf{p}_0)\ \mathrm{d}\mathbf{p}_0, \tag{S2}$$

$$p_{\mathrm{EF}}(\omega \mid \theta) = \int p(\omega \mid \theta, \mathbf{p}_0, \mathbf{u}_{\mathrm{EF}})\varphi(\mathbf{p}_0)\ \mathrm{d}\mathbf{p}_0, \tag{S3}$$

for trajectories simulated by the autonomous and electrotactic models, respectively. Each observed trajectory in the experimental training data set, $\mathbf{x}_{\mathrm{NoEF},i}$ and $\mathbf{x}_{\mathrm{EF},i}$, thus defines a set in the simulation space,

$$\Omega(\mathbf{x}_{\mathrm{NoEF},i}) = \left\{\omega = (\mathbf{x}(t), \mathbf{p}(t))_{t\geq 0} \ :\ \mathbf{x}(t_j) = \mathbf{x}_{\mathrm{NoEF},i}(t_j)\ \forall j = 0, \dots, 60\right\},$$

$$\Omega(\mathbf{x}_{\mathrm{EF},i}) = \left\{\omega = (\mathbf{x}(t), \mathbf{p}(t))_{t\geq 0} \ :\ \mathbf{x}(t_j) = \mathbf{x}_{\mathrm{EF},i}(t_j)\ \forall j = 0, \dots, 36\right\},$$

of all simulated trajectories that are indistinguishable from the observed training data. We thus define the likelihoods of each simulation as

$$\mathcal{L}_{\mathrm{NoEF}}(\mathbf{x}_{\mathrm{NoEF},i} \mid \theta) = \int_{\Omega(\mathbf{x}_{\mathrm{NoEF},i})} p_{\mathrm{NoEF}}(\omega \mid \theta)\ \mathrm{d}\omega,$$

$$\mathcal{L}_{\mathrm{EF}}(\mathbf{x}_{\mathrm{EF},i} \mid \theta) = \int_{\Omega(\mathbf{x}_{\mathrm{EF},i})} p_{\mathrm{EF}}(\omega \mid \theta)\ \mathrm{d}\omega,$$

for each cell index, $i$. The likelihoods of each trajectory thus combine to give the posterior,

$$\begin{aligned} \pi(\theta \mid \mathbf{x}_{\mathrm{NoEF}}, \mathbf{x}_{\mathrm{EF}}) &\propto \mathcal{L}(\mathbf{x}_{\mathrm{NoEF}}, \mathbf{x}_{\mathrm{EF}} \mid \theta)\pi(\theta) \\ &= \prod_i \mathcal{L}_{\mathrm{NoEF}}(\mathbf{x}_{\mathrm{NoEF},i} \mid \theta) \prod_j \mathcal{L}_{\mathrm{EF}}(\mathbf{x}_{\mathrm{EF},j} \mid \theta)\pi(\theta). \end{aligned} \tag{S4}$$

However, it is clear that the likelihood of each of the experimentally observed trajectories cannot easily be calculated. We therefore identified the posterior parameter distribution using a likelihood-free (i.e. simulation-based) Bayesian inference approach, harnessing the concept of *synthetic likelihoods*.

#### 3.1 Synthetic likelihoods

We focus on the autonomous case first; the electrotactic case follows similarly, with an obvious change of notation. The synthetic likelihood approach approximates the likelihoods, $\mathcal{L}_{\mathrm{NoEF}}(\mathbf{x}_{\mathrm{NoEF},i} \mid \theta)$ in two stages. The first stage is to reduce the dimension of the data space by defining a function of the simulated and observed trajectories that maps the data to a low-dimensional summary statistic. The second stage is to (a) use repeated simulation of the summarised model at the parameter value $\theta$ to fit an empirical multivariate Gaussian distribution for the summary statistic, and then (b) to approximate the likelihood with the *synthetic likelihood* of the experimental data, defined as the likelihood of the summarised data under the fitted empirical Gaussian distribution.

We define the function $Y : \omega \mapsto \mathbb{R}^4$ for the simulated trajectory $\omega = (\mathbf{x}(t), \mathbf{p}(t))$ on $t \in [t_n, t_{n+m}]$ as:

$$Y_1(\omega) = (\mathbf{x}(t_{n+m}) - \mathbf{x}(t_n)) \cdot \mathbf{i} \tag{S5a}$$

$$Y_2(\omega) = \|\mathbf{x}(t_{n+m}) - \mathbf{x}(t_n)\|, \tag{S5b}$$

$$Y_3(\omega) = \sum_{r=1}^{m} \|\mathbf{x}(t_{n+r}) - \mathbf{x}(t_{n+r-1})\|, \tag{S5c}$$

$$Y_4(\omega) = \left( \frac{1}{m} \sum_{r=1}^{m} \|\mathbf{x}(t_{n+r}) - \mathbf{x}(t_{n+r-1}) - Y_3/m\|^2 \right)^{1/2}, \tag{S5d}$$

for sample time points $t_j = 5j$ min. Thus, the entries of the vector $Y(\omega)$ denote the random values of

- the displacement over the interval $[t_n, t_{n+m}]$ in the positive $x$-direction,
- the net displacement,
- the path length,
- and the standard deviation of cell displacements over five-minute intervals,

for stochastic simulations $\omega$ of the electrotactic model in Eq. (3), given $\theta$, $\mathbf{p}_0$, and $\mathbf{u}(t)$. Note that we can also calculate the values of the function $Y$ in Eq. (S5) for the experimentally observed data, $\mathbf{x}_{\mathrm{NoEF},i}$, for each cell index, $i$. With a slight abuse of notation, we denote the summarised experimental data by $y_{\mathrm{NoEF},i} = Y(\mathbf{x}_{\mathrm{NoEF},i})$.

For a given value of $\theta$, the synthetic likelihood approach [21–23] assumes that the random value of $Y(\omega)$ under the density $p_{\mathrm{NoEF},i}(\omega \mid \theta)$ is a Gaussian random variable with parameter-dependent mean $\mu_{\mathrm{NoEF}}(\theta)$ and covariance $\Sigma_{\mathrm{NoEF}}(\theta)$. We estimate this mean and covariance with the sample mean and covariance of simulated summary statistics $Y(\omega_k)$, for $k = 1, \ldots, n$, produced by simulating the autonomous model $n$ times using the parameter value $\theta$ over the interval $t \in [0, 300]$. The resulting approximation of each trajectory's likelihood, $\tilde{\mathcal{L}}_{\mathrm{NoEF},n} \approx \mathcal{L}_{\mathrm{NoEF}}$, is summarised as

$$\tilde{\mathcal{L}}_{\mathrm{NoEF},n}(\mathbf{x}_{\mathrm{NoEF},i} \mid \theta) = \mathcal{N}\left(y_{\mathrm{NoEF},i} \mid \hat{\mu}_{\mathrm{NoEF}}(\theta), \hat{\Sigma}_{\mathrm{NoEF}}(\theta)\right) \quad i = 1, \ldots, 50, \tag{S6a}$$

$$\hat{\mu}_{\mathrm{NoEF}}(\theta) = \frac{1}{n} \sum_{k=1}^{n} Y(\omega_k), \tag{S6b}$$

$$\hat{\Sigma}_{\mathrm{NoEF}}(\theta) = \frac{1}{n} \sum_{k=1}^{n} (Y(\omega_k) - \hat{\mu}_{\mathrm{NoEF}}(\theta))(Y(\omega_k) - \hat{\mu}_{\mathrm{NoEF}}(\theta))^T, \tag{S6c}$$

$$\omega_k \sim p_{\mathrm{NoEF}}(\cdot \mid \theta) \quad k = 1, \ldots, n, \tag{S6d}$$

where $\mathcal{N}$ denotes the Gaussian density and where the chosen number of simulations, $n$, needs to be appropriately large [22]. In our case, we choose $n = 500$.

In the case of the electrotactic model with piecewise constant electric field input, $\mathbf{u}_{\mathrm{EF}}(t)$, we adapt the procedure above to summarise the simulations and the training data with an eight-dimensional summary statistic. Treating the intervals $t \in [0, 60], [60, 180], [180, 300]$ separately, we summarise simulations, $\omega \sim p_{\mathrm{NoEF}}(\cdot \mid \theta)$, and observations, $\mathbf{x}_{\mathrm{EF},i}$, by calculating $Y$ for each interval. We use only $t \leq 180$ to calibrate the model, while holding back the interval $t \in [180, 360]$ for the purpose of testing. Thus, we combine the values of $Y$ for the intervals $t \in [0, 60]$ and $t \in [60, 180]$ into an eight-dimensional summary for calibrating the model. Using this eight-dimensional summary, we adapt the synthetic likelihood procedure summarised in Eq. (S6) to calculate $\tilde{\mathcal{L}}_{\mathrm{EF},n}(\mathbf{x}_{\mathrm{EF},i} \mid \theta)$.

Finally, we can multiply each of these trajectory synthetic likelihoods into an overall synthetic likelihood for the experimental data,

$$L_{\mathrm{NoEF},n}(\theta) = \prod_i \tilde{\mathcal{L}}_{\mathrm{NoEF},n}(\mathbf{x}_{\mathrm{NoEF},i} \mid \theta), \tag{S7a}$$

$$L_{\mathrm{EF},n}(\theta) = \prod_i \tilde{\mathcal{L}}_{\mathrm{EF},n}(\mathbf{x}_{\mathrm{EF},i} \mid \theta), \tag{S7b}$$

$$\mathcal{L}(\mathbf{x}_{\mathrm{NoEF}}, \mathbf{x}_{\mathrm{EF}} \mid \theta) \approx L_n(\theta) = L_{\mathrm{NoEF},n}(\theta) L_{\mathrm{EF},n}(\theta), \tag{S7c}$$

each calculation of which requires $n$ simulations of the autonomous model and $n$ of the the electrotactic model.

#### 3.2 SMC inference

In order to produce a sample from the posterior distribution, we use sequential Monte Carlo (SMC) with synthetic likelihoods [21–27], as outlined in Algorithm 1. This method is chosen in order to exploit parallelisation, mitigating the computational burden of MCMC-based approaches that is incurred due to the large numbers of model simulations required for accurate likelihood-free inference. SMC defines a sequence of intermediate importance distributions that evolve towards the target posterior. This approach is particularly useful in comparison to naive rejection sampling: since we will use non-informative priors, rejection sampling is too inefficient, as it proposes parameters in extremely low-likelihood regions of parameter space too frequently. Importantly, for each value of $\theta$, the stochastic values of $L_{\mathrm{NoEF},n}(\theta)$ and $L_{\mathrm{EF},n}(\theta)$ can be called multiple times. At each call of these two likelihoods, we do *not* recycle previously computed values for the synthetic likelihoods but instead simulate the models again. Although this approach slows the inference procedure, it is necessary to ensure the correct stationary distribution of the Markov chain steps, (13–14).

In Algorithm 1 we produce a weighted sample from the Bayesian synthetic likelihood approximation to the posterior, $\pi(\theta \mid \mathbf{x}_{\mathrm{NoEF}}, \mathbf{x}_{\mathrm{EF}})$. The intermediate distributions at each iteration are proportional to the tempered distributions

$$\pi_T(\theta) \propto \left[L_{\mathrm{NoEF},n}(\theta) L_{\mathrm{EF},n}(\theta)\right]^T \pi(\theta),$$

where the sequence of temperatures $T$ evolves from 0 to 1. In Algorithm 1, we define the initial perturbation kernel, $K(\cdot \mid \theta)$ to be a multivariate Gaussian density with mean $\theta$ and diagonal covariance matrix, with component-wise variances of 0.01, 0.0025 and 0.0001 for $v$, $\Delta W$, and $D$, respectively, and 0.01 for each $\gamma_i$. The effective sample size, $ESS$, used to adaptively choose the increment in temperature, is defined as

$$ESS(\{W_i\}) = \left(\sum_i W_i\right)^2 / \sum_j W_j^2, \tag{S8}$$

for any finite set of sample weights, $W_i$. We use $N = 1000$ particles, setting $N_{\min} = 333$ as the effective sample size triggering resampling, and setting $\alpha = 0.8$ as the decay rate of the effective sample size.

The effective sample size of the sample produced by Algorithm 1, as defined in Equation (S8), can be an overestimate of the sample quality. The resampling step, in lines (9–11), tends to result in multiple particles with equal parameter values. These particles do not always separate through the single MCMC step in (13–14). The replicated parameter values in the sample thus degrade its quality, without being captured by the ESS calculation. To improve the quality of the final sample produced by Algorithm 1, we post-process the posterior. First, the final weighted Monte Carlo sample output from Algorithm 1 is resampled according to steps (9–11). Then, for each resampled particle, we calculate 100 MCMC steps (13–14) using $T = 1$. This procedure effectively produces 1000 short Markov chains of length 100, each beginning from a particle from the target distribution, and with stationary distribution equal to the target distribution. We use the end samples of each of these Markov chains as

**Algorithm 1** Synthetic Likelihood SMC

**Input:** Observed summary statistics $y_{\mathrm{NoEF}}$ and $y_{\mathrm{EF}}$; prior $\pi$; perturbation kernel $K(\cdot \mid \theta)$; .

**Output:** Weighted sample set of parameters $\theta_i$ with weights $W_i$, from the synthetic likelihood approximation to the posterior $\pi(\theta \mid \mathbf{x}_{\mathrm{NoEF}}, \mathbf{x}_{\mathrm{EF}})$.

Sample $N$ independent $\theta_i$ from $\pi$.

Set weights $W_i^0 = 1/N$ for $i = 1, \ldots, N$.

Initialise $T = 0$ and $r = 0$.

**repeat**

  Update $r \leftarrow r + 1$.

  Find $\Delta T \in [\Delta T_{\min}, 1 - T]$ to solve $ESS(\{W_i^r\}) = \alpha ESS(\{W_i^{r-1}\})$, for weights $W_i^r$ such that

$$\log W_i^r = \log W_i^{r-1} + \Delta T \left( \log L_{\mathrm{NoEF},n}(\theta_i) + \log L_{\mathrm{EF},n}(\theta_i) \right),$$

  for the synthetic likelihoods, $L_{\mathrm{NoEF},n}(\theta_i)$ and $L_{\mathrm{EF},n}(\theta_i)$. Use $\Delta T = \Delta T_{\min}$ or $\Delta T = 1 - T$ if $ESS(\{W_i^r\})$ is, respectively, uniformly less than or uniformly greater than $\alpha ESS(\{W_i^{r-1}\})$ on the interval $[\Delta T_{\min}, 1 - T]$.

  Update $T \leftarrow T + \Delta T$.

  **if** $ESS(\{W_i^r\}) < N_{\min}$ **then**

    Resample from $\{\theta_i\}$ according to weights $W_i^r$.

    Reset weights $W_i^r = 1/N$.

    Reset covariance of perturbation kernel, $K$, to equal the empirical covariance of the resampled parameter set, $\{\theta_i\}$.

  **end if**

  Propose perturbed parameter values $\theta_i^\star \sim K(\cdot \mid \theta_i)$ for $i = 1, \ldots, N$.

  Accept $\theta_i \leftarrow \theta_i^\star$ with Metropolis–Hastings probability

$$\left[ \frac{L_{\mathrm{NoEF},n}(\theta_i^\star) L_{\mathrm{EF},n}(\theta_i^\star)}{L_{\mathrm{NoEF},n}(\theta_i) L_{\mathrm{EF},n}(\theta_i)} \right]^T \frac{\pi(\theta_i^\star) K(\theta_i^\star \mid \theta_i)}{\pi(\theta_i) K(\theta_i \mid \theta_i^\star)}$$

  for $i = 1, \ldots, N$.

**until** $T = 1$.

a higher-quality, 1000-particle sample from the Bayesian synthetic likelihood approximation to the posterior, each with equal weight. The resulting particles are checked to represent distinct parameter values, and thus every sample generated is of size 1000, where 'size' refers to both particle numbers and effective sample size, which are identical.

### 4 Model validation

#### 4.1 Autonomous data cross-validation

We first evaluate the modelling approach by comparing the model outputs trained to each of the two replicates alone in turn, with the other replicate held back. Note that to produce a posterior based on the autonomous (control) data set alone, we can simply adapt Algorithm 1 to use only $L_{\mathrm{NoEF},n}$ in calculating synthetic likelihoods (effectively setting $L_{\mathrm{EF},n} = 1$). The one-dimensional marginals of the resulting posterior, trained on the entire autonomous data set, are shown in Figure 2 of the main text. The covariance structure of the posterior sample is depicted in Figure S2.

If training on each of the two control data sets alone, we can construct an additional two posteriors. Figure S3 demonstrates that these posteriors closely overlap with one another, and with the posterior trained on the combined data set. In addition to comparing the posteriors produced by each replicate, we can use repeated simulation to produce posterior predictive distributions for each of the four summary statistics used for inference. These can be compared to each of the observed data sets. In Figure S4, we plot posterior predictive distributions, based on 10 simulations for each of the 1000 sampled parameter values, for each of the three posteriors. In this figure, we demonstrate that there is a good agreement between these posterior predictive distributions and the empirically observed summary statistics for each replicate of the autonomous experiment.

We can quantify this agreement by estimating the log-likelihood of each data set, using the maximum likelihood normal approximations to each of the three empirical distributions depicted in Figure S4. The quantified cross-validation is shown in Table S1. For the posterior trained on replicate 1, the posterior predictive log-likelihood of the data from replicate 1 is, as expected, slightly greater than that of replicate 2. However, the drop in log-likelihood for the replicate 2 data is minimal, and thus the out-of-sample predictive performance of the posterior trained against replicate 1 is good. A similar cross-validation argument, swapping the roles of replicate 1 and replicate 2, holds. Thus, Figure S4 and Table S1 validate the suitability of the calibrated autonomous model.

Finally, the preliminary analysis of the observed displacements in each replicate, depicted in Figure S1, can be replicated for simulated trajectories using parameters sampled from the posteriors trained on each replicate alone. Figure S5 shows that model simulations produce qualitatively similar behaviour to the observed data. Importantly, these characteristics of the trajectories were not explicitly used to calibrate the model. However, they have been replicated by the calibrated model, which provides further evidence that the autonomous model is accurate.

#### 4.2 Electrotactic model validation

After validating the autonomous model against the control data, we perform the synthetic likelihood inference procedure on the full training data set, comprising each observed trajectory from both control data sets, $\mathbf{x}_{\mathrm{NoEF},i}$ on $t \in [0, 300]$, and the first section of each observed trajectory from the electrotactic data sets, $\mathbf{x}_{\mathrm{EF},i}$ on $t \in [0, 180]$. Algorithm 1 is used to construct 16 separate posteriors, based on each of the 16 priors, $\pi_X$, indexed by $X \subset \{1, 2, 3, 4\}$, where $\gamma_i > 0$ if and only if $i \in X$. The greatest posterior likelihood is given by choosing $X = \{4\}$, and thus setting $\gamma_1 = \gamma_2 = \gamma_3 = 0$. For this choice of prior, we plot the one-dimensional marginals of the resulting posterior in Figure 4 of the main text. To illustrate the covariance structure of the resulting posterior, the two-dimensional marginals are given in Figure S6.

To validate the resulting posterior, in Figure S7 we compare the one-dimensional marginals of the parameters $v$, $\Delta W$, and $D$ for the posteriors trained against the autonomous data set only, versus the posterior trained against the entire training data set. The resulting posteriors are similar, showing that calibrating to the full training data set refines the predictions of model calibrated to the autonomous data set alone.

Moreover, the posterior predictive distributions of the four summary statistics over 0 min to 300 min (for the autonomous experiment) and each of 0 min to 60 min and 60 min to 180 min (for the electrotactic experiment), depicted in Figure S8, show that the model is a close fit to the observed summarised trajectories in the training data set. In particular, similarly to the autonomous model above, we have split the training data between replicates 1 and 2, and trained posteriors on each. We can perform cross-validation analysis, using the quantification of the log-likelihood of the observed summary statistics according to the posterior log-likelihoods in Table S2. As with the autonomous model, this table shows that the posteriors trained on each replicate provide a good prediction of the log-likelihood of the summary statistics of the other replicate. Thus, the calibrated model provides good out-of-sample predictive performance.

Note that, in addition to this cross-validation approach, a further test of the model validity is depicted in Figure 5 of the main manuscript. We use the electrotactic model, calibrated to the training data, $\mathbf{x}_{\mathrm{NoEF}}$ and $\mathbf{x}_{\mathrm{EF}}$, to predict the behaviour of the held-back test data set, $\mathbf{x}_{\mathrm{Switch}}$, comprising the observed cell trajectories over $t \in [180, 360]$ after the EF switches direction from the positive to negative $x$-direction. We demonstrate that the posterior predictive distributions of the summary statistics are a good match to the unseen test data, further confirming the ability of the calibrated model to predict cellular motility under dynamic EF inputs.

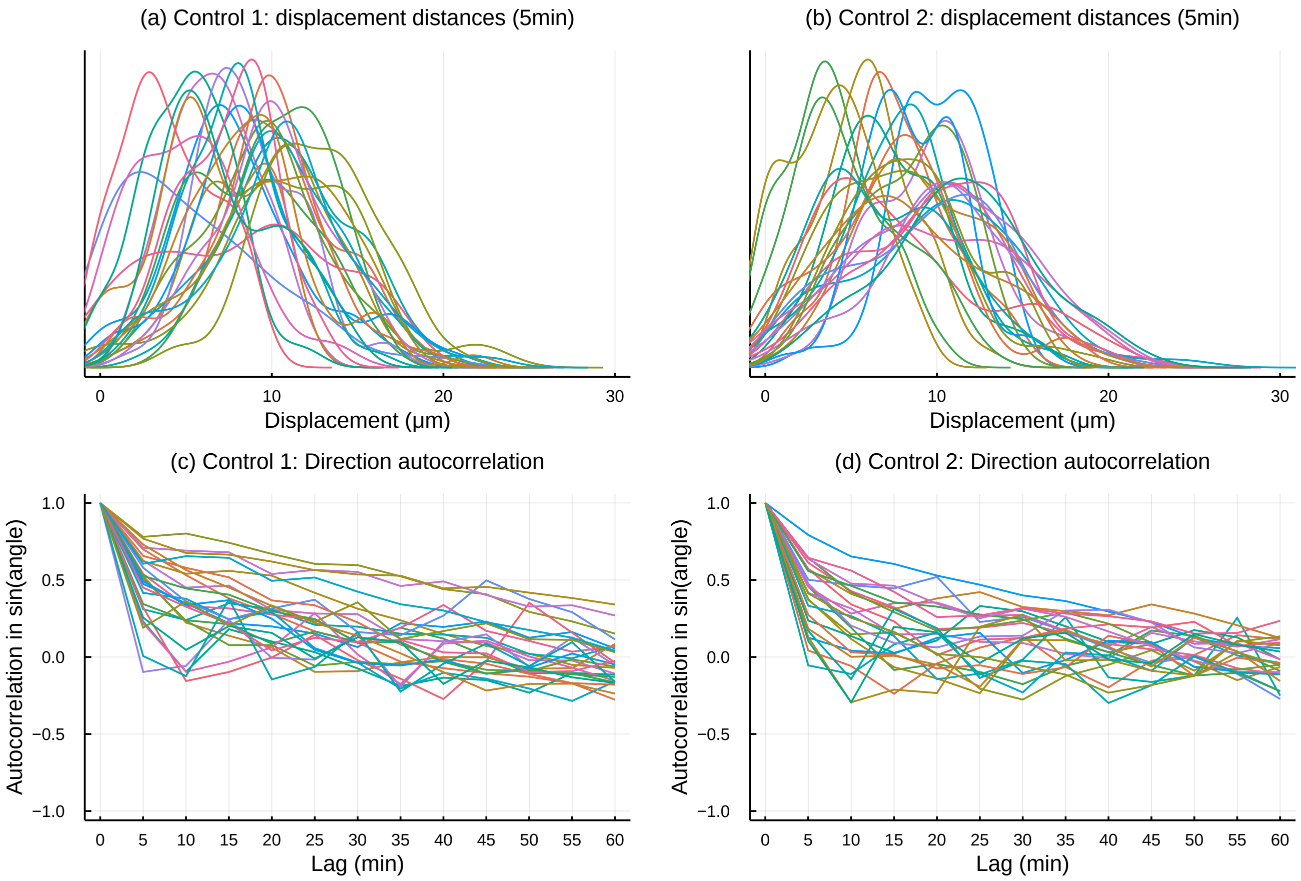

Figure S1: Preliminary analysis of autonomous data sets. (a, b) Distributions of observed displacement distances for each tracked cell in each autonomous data set. (c, d) Autocorrelations of observed displacement angles for all tracked cells in each autonomous data set, for intervals at lags of 5 min to 60 min.

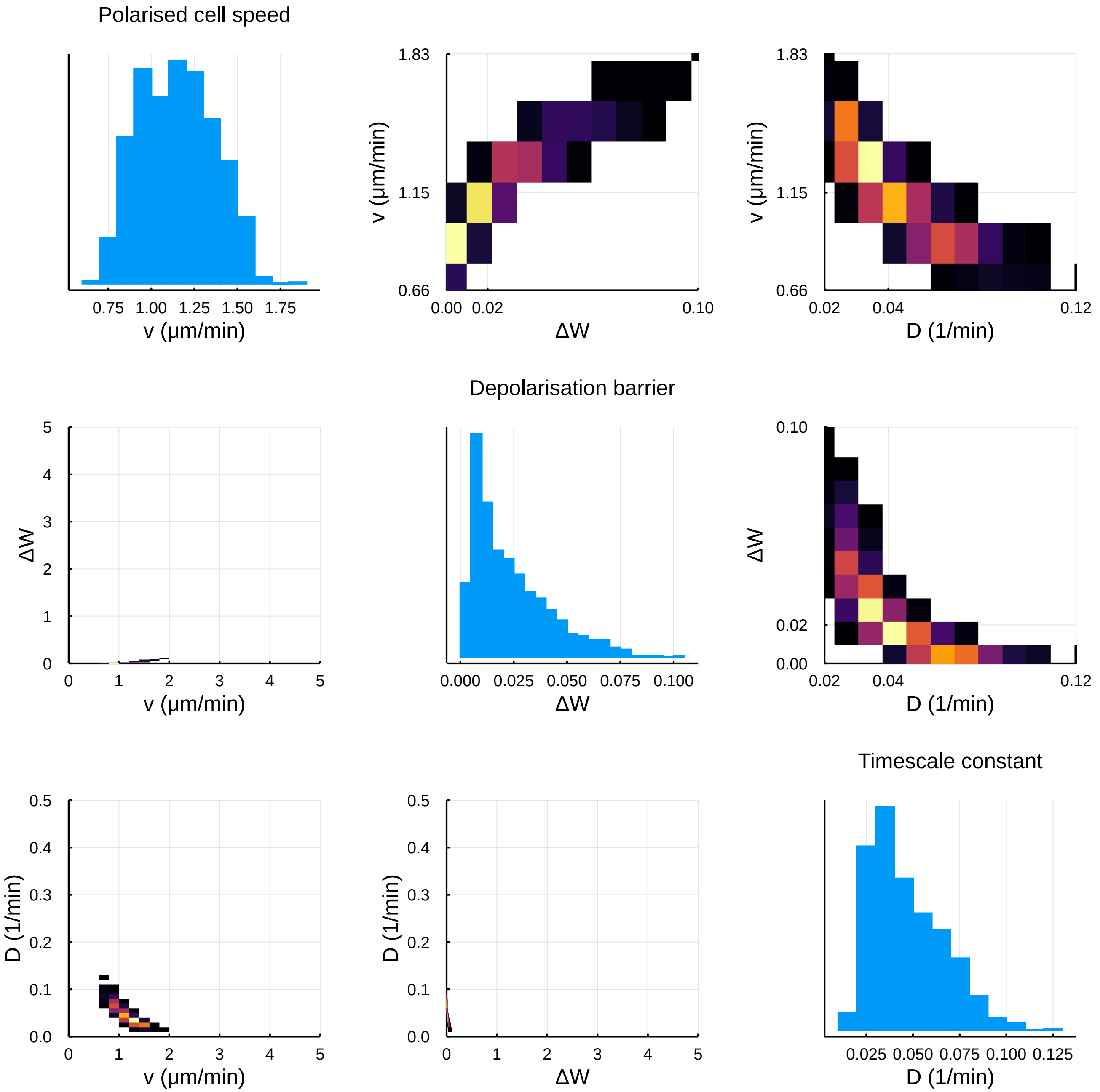

Figure S2: Posterior sample from $\pi(\theta \mid \mathbf{x}_{\mathrm{NoEF}})$, generated by Algorithm 1 neglecting $L_{\mathrm{EF},n}$, for identified parameters, $v$, $\Delta W$, and $D$. Diagonal plots are empirical histograms (as in Figure 2 in the main text). Off-diagonal heatmaps represent empirical pairwise distributions, where brighter colours correspond to greater density. Axes in the bottom-left are scaled to the prior support; the top-right scales are zoomed in to the region of greatest positive posterior likelihood.

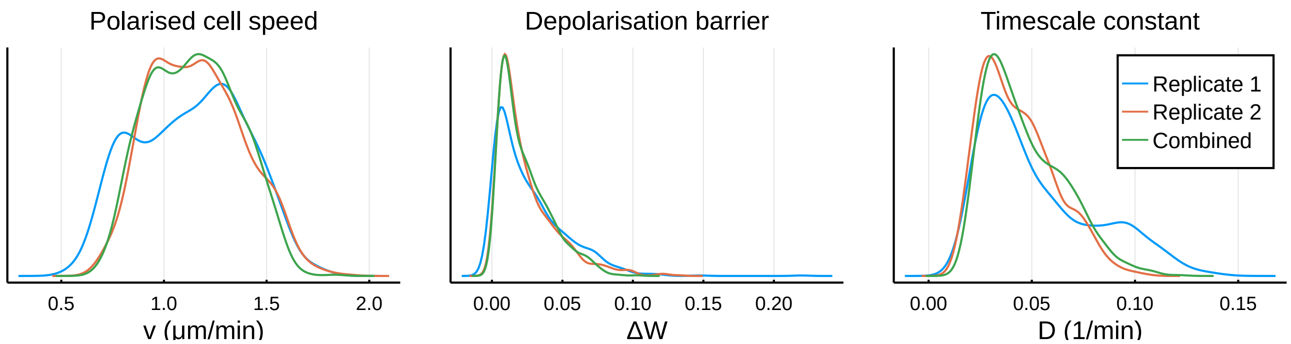

Figure S3: Posterior distributions for the parameters of the autonomous model, trained to each replicate, and also to the combined data set. Posterior samples have been generated using Algorithm 1, with MCMC post-processing, and depicted as densities using a default KDE.

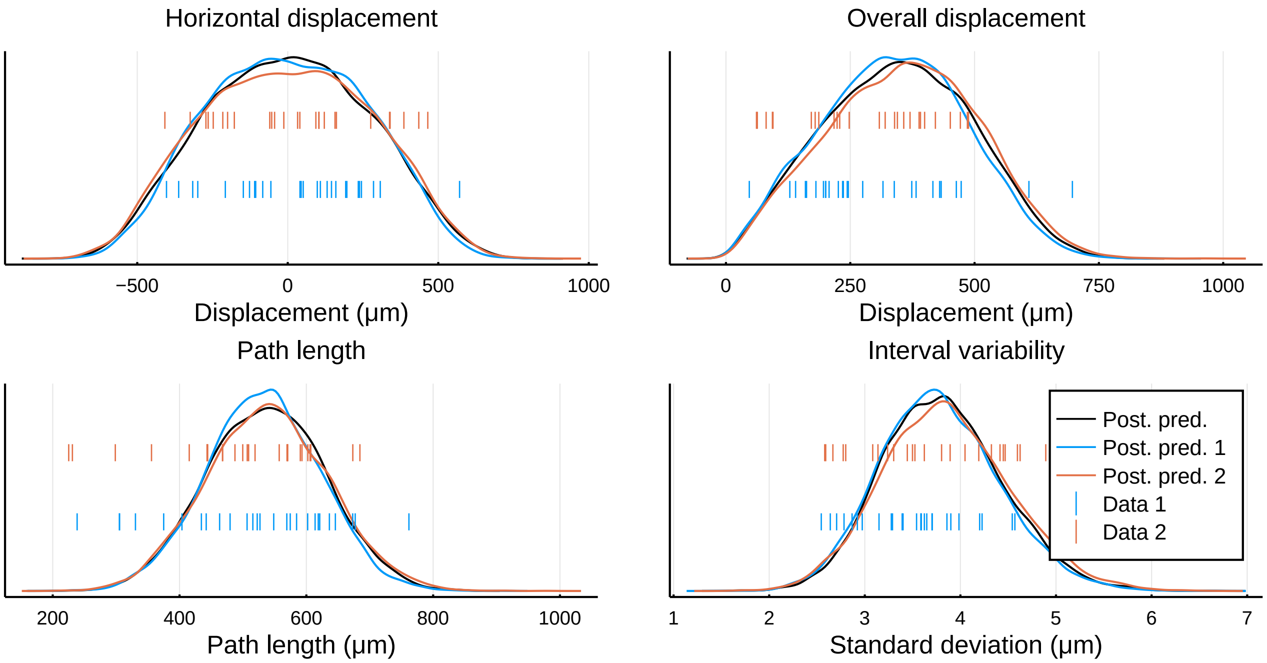

Figure S4: Posterior predictive distributions of the summary statistics used for inference, for autonomous data $\mathbf{x}_{\mathrm{NoEF}}$ only. For each posterior sample trained against replicate 1, replicate 2, and both replicates together, we simulated 10,000 summary statistics to produce three posterior predictive distributions (kernel density estimates, represented as solid curves). These are plotted with the observed summary statistics from each of the data sets. The posterior predictive distributions for each replicate can thus be cross-validated against the data in the other replicate: the log-likelihoods of the observations in each replicate, under each posterior predictive distribution, are recorded in Table S1.

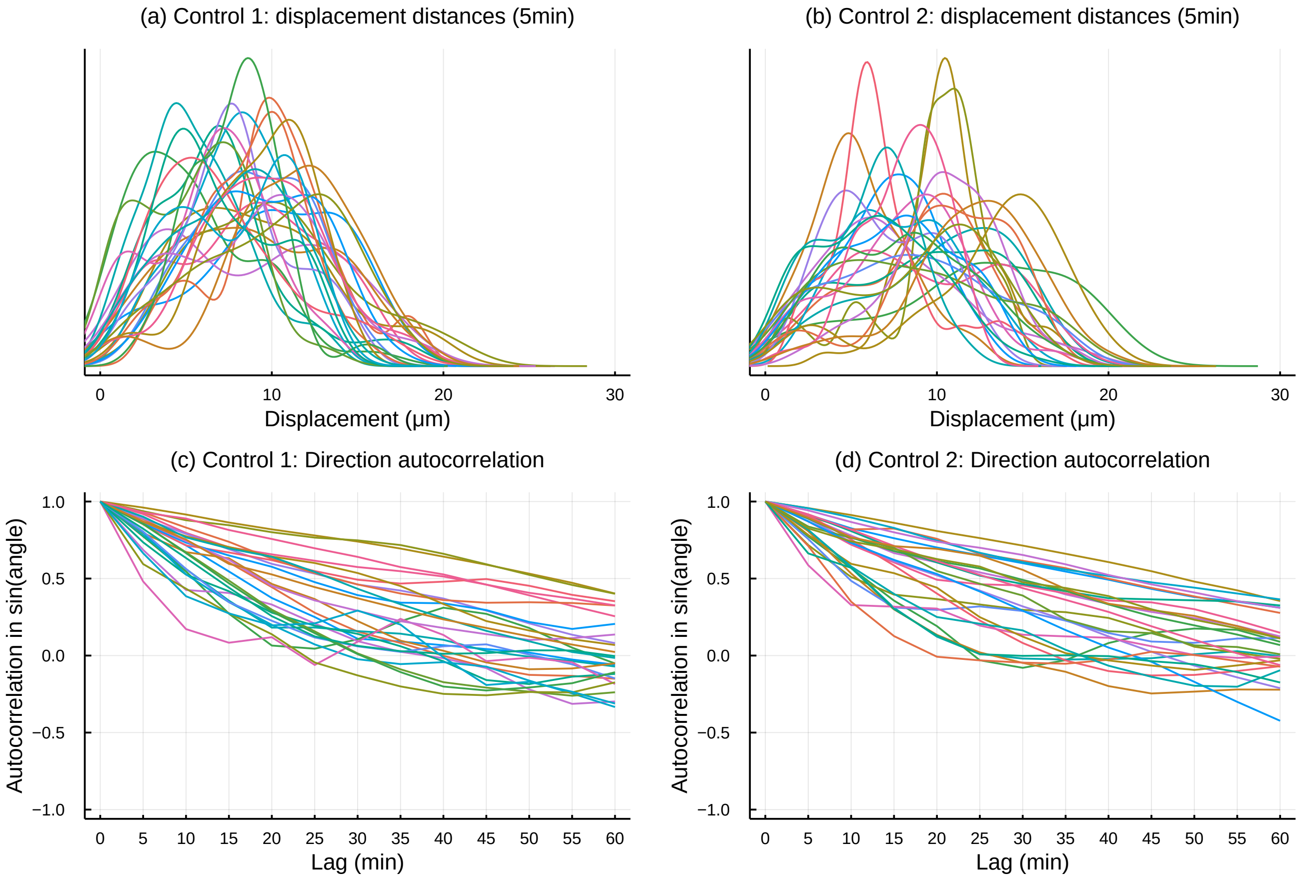

Figure S5: Analysis of simulated displacements over 5 min intervals. (a, b) Distributions of simulated displacement distances for cells simulated using parameters sampled from posteriors trained on each autonomous data set. (c, d) Autocorrelations of simulated displacement angles for cells simulated using parameters sampled from posteriors trained on each autonomous data set, for intervals at lags of 5 min to 60 min.

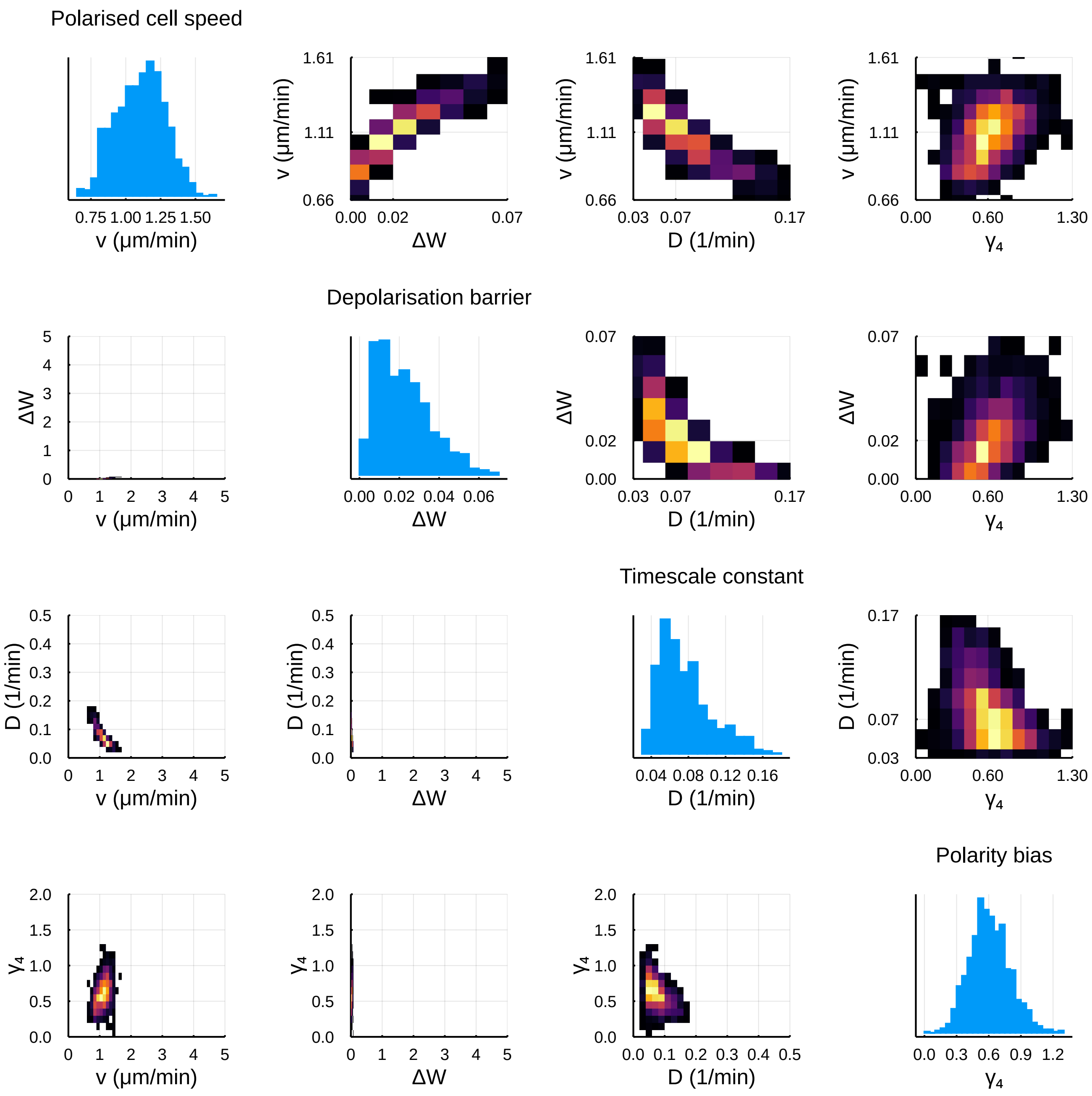

Figure S6: Weighted sample from $\pi(\theta \mid \mathbf{x}_{\mathrm{NoEF}}, \mathbf{x}_{\mathrm{EF}})$, generated by the completion of Algorithm 1 for identified parameters, $v$, $\Delta W$, $D$, and $\gamma_4$. Plots are as described in Figure S2.

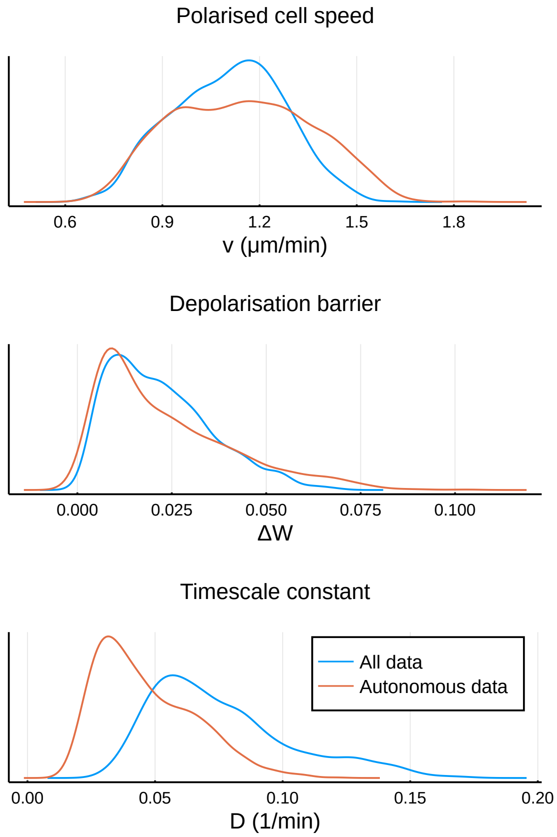

Figure S7: One-dimensional projections for $v$, $\Delta W$, and $D$ of the empirical posteriors $\pi(\theta \mid \mathbf{x}_{\mathrm{NoEF}}, \mathbf{x}_{\mathrm{EF}})$ and $\pi(\theta \mid \mathbf{x}_{\mathrm{NoEF}})$, generated by Algorithm 1, trained on all data and the autonomous data, respectively.

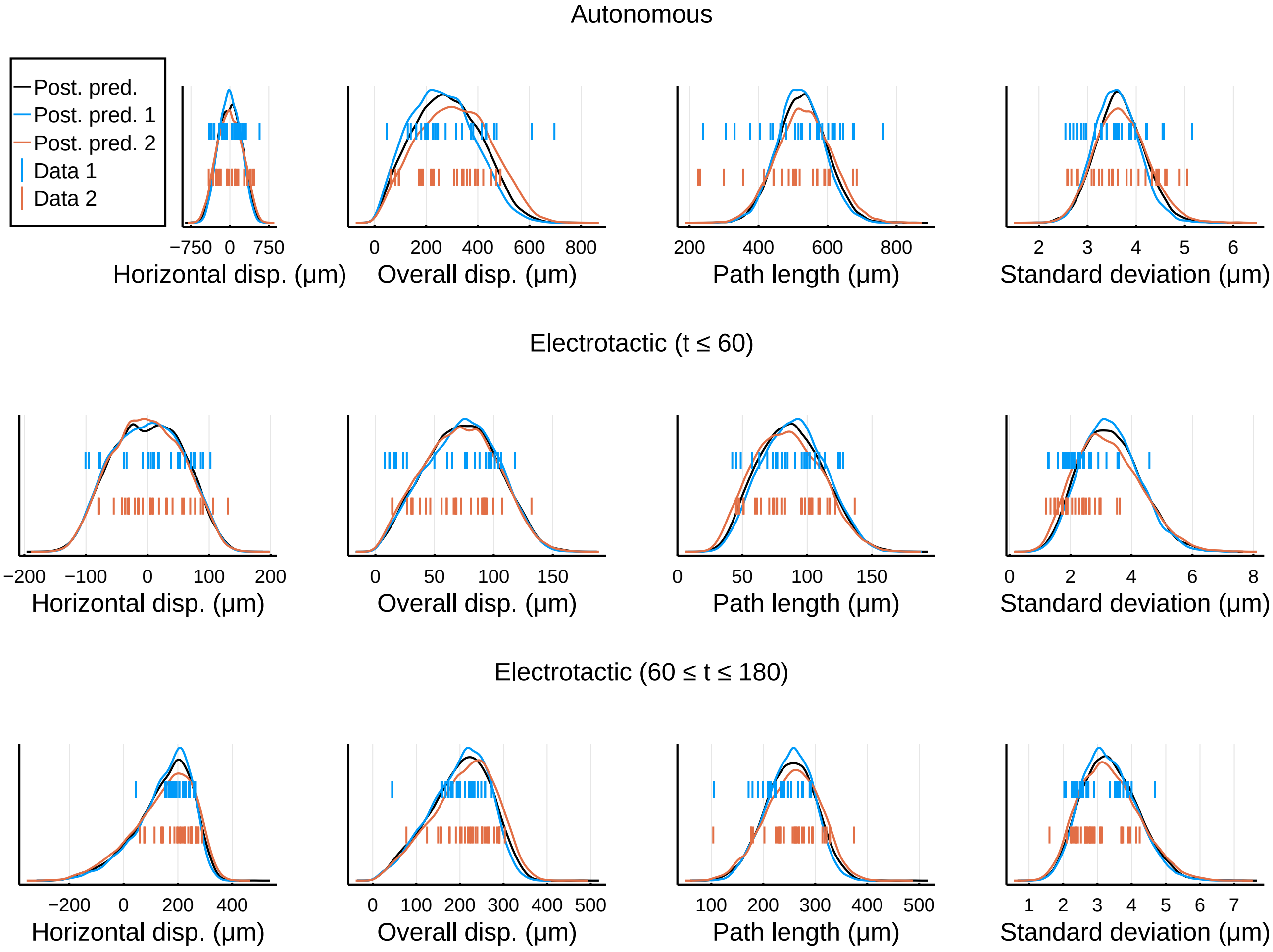

Figure S8: Posterior predictive distributions of all summary statistics used for inference with combined training data $\mathbf{x}_{\mathrm{NoEF}}$ and $\mathbf{x}_{\mathrm{EF}}$. Details are as for Figure S4, with log-likelihoods of the observations under each posterior predictive distribution recorded in Table S2.

### List of Tables

S1 Cross-validated log-likelihoods of training data under empirical posterior predictive distributions: autonomous data only. . . . . 19
S2 Cross-validated log-likelihoods of training data under empirical posterior predictive distributions: all training data. . . . . 20

| Log likelihoods | Replicate 1 training data:<br>27 autonomous trajectories | Replicate 2 training data:<br>26 autonomous trajectories |
|---|---|---|
| Replicate 1 posterior | −558.6 | −537.4 |
| Replicate 2 posterior | −559.9 | −537.2 |
| Combined posterior | −559.3 | −537.5 |

Table S1: Cross-validated log-likelihoods of training data under empirical posterior predictive distributions, trained on data from the autonomous experiment only. Entries $(i, j)$ correspond to the log-likelihood of observed data from Replicate $j$ under the posterior predictive distribution for posteriors trained on data from Replicate $i$, for $i, j \in \{1, 2\}$. The bottom row corresponds to log-likelihoods under the posterior predictive distribution for the posterior trained on the combined data set. This table quantifies the log-likelihood of Figure S4.

| Log likelihoods | Replicate 1 training data:<br>27 autonomous trajectories<br>26 electrotactic trajectories | Replicate 2 training data:<br>26 autonomous trajectories<br>30 electrotactic trajectories |
|---|---|---|
| Replicate 1 posterior | −1468 | −1550 |
| Replicate 2 posterior | −1497 | −1543 |
| Combined posterior | −1479 | −1547 |

Table S2: Cross-validated log-likelihoods of training data under empirical posterior predictive distributions, trained on all identified training data. Details are as for Table S1. This table quantifies the log-likelihood of Figure S8.
